# Three-dimensional Monolayer Stress Microscopy

**DOI:** 10.1101/616987

**Authors:** Ricardo Serrano, Aereas Aung, Yi-Ting Yeh, Shyni Varghese, Juan C. Lasheras, Juan C. del Álamo

**Affiliations:** University of California San Diego; Massachusetts Institute of Technology; Duke University

## Abstract

Many biological processes involve the collective generation and transmission of mechanical stresses across cell monolayers. In these processes, the monolayer undergoes lateral deformation and bending due to the tangential and normal components of the cell-generated stresses. Monolayer Stress Microscopy (MSM) methods have been developed to measure the intracellular stress distribution in cell monolayers. However, current methods assume plane monolayer geometry and neglect the contribution of bending to the intracellular stresses.

This work introduces a three-dimensional (3D) MSM method that calculates monolayer stress from measurements of the 3D traction stresses exerted by the cells on a flexible substrate. The calculation is carried out by imposing equilibrium of forces and moments in the monolayer, subject to external loads given by the 3D traction stresses. The equilibrium equations are solved numerically, and the algorithm is validated for synthetic loads with known analytical solutions.

We present 3D-MSM measurements of monolayer stress in micropatterned islands of endothelial cells of different sizes and shapes. These data indicate that intracellular stresses caused by lateral deformation emerge collectively over long distances; they increase with the distance from the island edge until they reach a constant value that is independent of island size. On the other hand, bending-induced intracellular stresses are more concentrated spatially and remain confined to within 1-2 cell lengths of bending sites. The magnitude of these bending stresses is highest at the edges of the cell islands, where they can exceed the intracellular stresses caused by lateral deformations. Our data from non-patterned monolayers suggests that biomechanical perturbations far away from monolayer edges also cause significant localized alterations in bending tension. The localized effect of bending-induced stresses may be important in processes like cellular extravasation, which are accompanied by significant normal deflections of a cell monolayer (i.e. the endothelium), and which require localized changes in monolayer permeability.

## 1 Introduction

Cells interact with their environment via biochemical and biophysical signals, including mechanical signals mediated by contractile forces^1^. When a cell contracts, it builds up intracellular stresses that can be transmitted to the extracellular matrix and to adjacent cells. This collective generation and transmission of intracellular stress in thin multicellular colonies plays an important role in biological processes such as development^2^, endothelial function^3^ and wound healing^4^. The development of microscopy methods for the measurement of intracellular stresses has improved our fundamental understanding of these biological processes^5^. Furthermore, the high-throughput quantification of intracellular stresses provides great promise in the functional screening of contractile tissues for drug development^6, 7^.

Monolayer Stress Microscopy (MSM) quantifies the spatial distribution of intracellular stress in confluent few-cell-thick cultures^6, 8-12^. Most MSM methods rely on *a priori* knowledge of the traction stresses that the cells exert on their substrate, which are conceptualized as loads acting on the monolayer by Newton’s third law^8, 9^. Several alternative models have been proposed to formulate the calculation of intracellular stresses in closed mathematical form. Pioneering MSM methods prescribed a compatibility condition consistent with a linearly elastic constitutive relation for the monolayer, as well as the boundary loads at the edges of the domain^8^. Subsequent efforts^10^ have modeled cells in the monolayer as a set of particles that can exert forces on other particles according to a potential, and have used Hardy’s stress formula^13^ to infer the continuum stress distribution in the monolayer from the inter-particle forces. More recently, Bayesian inversion has been proposed to effectively replace explicit compatibility and boundary conditions by a set of sound assumptions about the symmetry and statistical noise of the intracellular stress tensor^12^. These methods have been shown to yield results that are generally in good agreement, but they all neglect the contribution of bending moments and normal traction stresses.

Even if they form thin layers, endothelia and epithelia often generate or are subjected to three-dimensional forces that cause monolayer bending. Epithelial bending and invagination are a key part of morphogenesis^14^. Besides, endothelial transmigration is accompanied by localized endothelial bending ^15^. Conceptually, a cell monolayer subjected to three-dimensional external loads can undergo bending due to the normal (i.e. out-of-plane) loads in addition to the lateral deformation caused by tangential (i.e. in-plane) loads (Figure 1A-C)^15, 16^. In principle, both the bending and the lateral deformations can cause intracellular stress (Figure 1D). Existing MSM methods assume that the contribution to intracellular stress from bending and normal traction stresses is negligible but this assumption contrasts with experimental evidence that cell monolayers generate significant normal traction stresses^9, 17^. The relative contribution of bending to the total intracellular stress in cell monolayers has not been measured experimentally yet.

**Figure 1.**
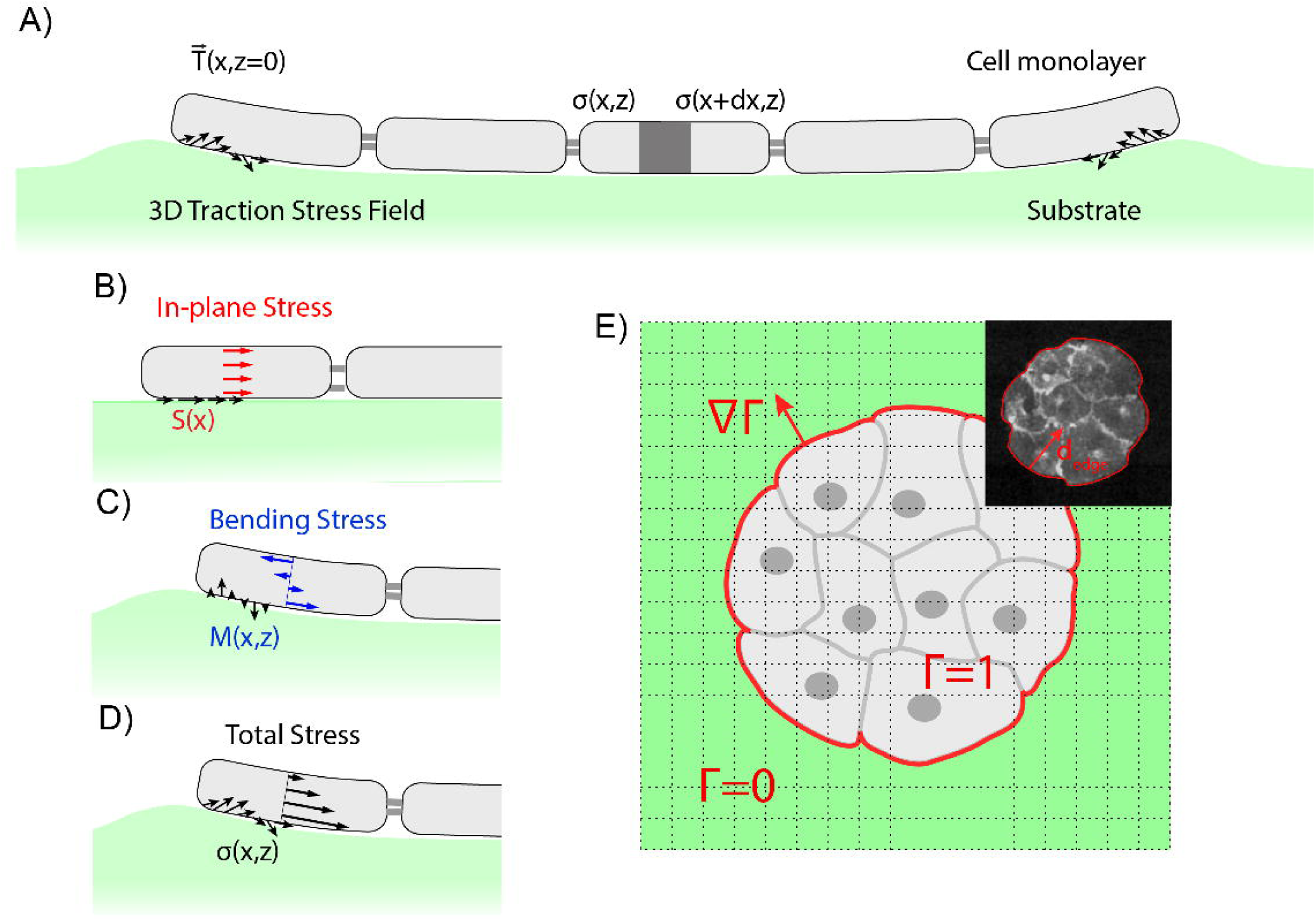
(A) Sketch of a cell monolayer exerting 3D traction stresses (arrows) on a deformable substrate. (B) Distribution of lateral stresses resulting from the tangential component of traction stress. (C) Distribution of bending stresses resulting from the normal component of traction stress. (D) The monolayer stress field is a combination of both lateral and bending effects. (E) Representation of a cell island (gray) in a Cartesian grid. The red line represents the edge of the monolayer that separates the interior of the island *Γ* = 1, from the outside *Γ* = 0. *∇ Γ*, is defined as a unitary normal vector pointing outside of the cell domain. Inset: example of the specified outline (red) in an experiment; *d*_*edge*_ is the distance from the edge of the monolayer.

This paper has two main parts. The first part introduces a new MSM method that accounts for bending effects to derive the intracellular stresses in a cell monolayer from the three-dimensional (3D) traction stresses generated by the monolayer. In the second part, this new technique is applied experimentally to multicellular micro patterned islands of different sizes and shapes, and to non-patterned monolayers subjected to a localized biomechanical perturbation.

Three-dimensional MSM infers the intracellular stresses caused by lateral deformation from the tangential traction stresses exerted by the cells on their substrate, while the intracellular stresses caused by monolayer bending are obtained from the normal traction stresses. The problem is formulated mathematically as a set of partial differential equations by modeling the cell monolayer as a thin linearly elastic plate, and by imposing equilibrium of forces and moments. The numerical integration of the equilibrium equations is carried out using a Fourier-Galerkin method, which allows for seamless integration with the 3D Fourier traction microscopy method used to measure the traction stresses at the interface between the cells and the substrate^18, 19^. An immersed interface method is used to apply the boundary conditions at the edges of the cell islands, which are defined implicitly using a level set function ^20^. This numerical integration procedure is validated using the known analytical solutions of a family of synthetic deformation fields that is representative of our experimental measurements.

Our experimental results indicate that lateral monolayer stresses emerge collectively over long distances consistent with previous observations ^21^. They increase towards the inner side of the island until they reach a plateau value that is independent of island size. On the other hand, bending-induced intracellular stresses are more concentrated spatially, and co-localize with sites of high normal deformation of the monolayer. The bending-induced intracellular stresses are particularly large at the edges of the cell islands or at sites where the monolayer is perturbed externally, where they can exceed the intracellular stresses caused by lateral deformations. The effect of the localization of bending stresses might be relevant in processes like neutrophil extravasation or cancer cell invasion where a cell disrupts an endothelial monolayer locally by exerting forces that are normal to the monolayer and its substrate ^15^ ^22^ ^23^. While the transmission of forces across cell junctions is recognized to be three-dimensional even in cell monolayers, the quantification of these forces has been mostly limited to two-dimensions so far. The present study provides a novel tool to quantify the three-dimensional transmission of forces across cell monolayers.

## 2 Materials and Methods

### 2.1 Micropatterned Polyacrylamide Gel Preparation

Square #1 glass coverslips of 22-mm in size were activated with a drop of 0.1M NaOH on a plate at 90 °C until drop evaporation. The coverslips were then washed with distilled water, dried and treated with 3-aminopropyl-trimethoxysilane for 3 min. These coverslips were rinsed with distilled water, dried and treated with 0.5% glutaraldehyde for 30 min. The activated surfaces were rinsed with distilled water and kept at room temperature for their use within the same day.

Round 12-mm glass coverslips were pretreated in a UV Ozone (UVO) box for 5 min. The coverslips were then incubated with a 110 μL drop of 0.2 mg/mL PLL-PEG for 30 min at room temperature. The chrome face of the photomask (Advance Reproductions, North Andover, MA) was activated by UVO for 3 min. The PLL-PEG coated coverslips were attached to the activated side of the photomask by sandwiching a 2.5 μL drop of distilled water between both surfaces. The assembly of the photomask and coverslips was then exposed to UVO light for 7 min. The photomask was detached from the coverslips by adding distilled water and then dried by aspirating excess water. A 110 μL drop of 50 μg/mL fibronectin (FN, Sigma-Aldrich, St. Louis, MO) was placed on the PLL-PEG coated surface of the coverslip and incubated for 40 mins. A thin film of FN was deposited on the areas of the coverslips that had been exposed to UVO light, whereas PLL-PEG prevented FN adhesion to the glass surface in the shadowed areas^24^.

The polyacrylamide gels were fabricated with a mixture of acrylamide and bis-acrylamide following a well-established method^9, 19, 25-30^. Fluorescent 0.2-μm diameter beads were added to the mixture for later use as fiduciary markers of gel deformation. To promote bead distribution towards the surface of the gel, phosphate-buffered saline (PBS) was used instead of distilled water. The Young’s modulus was controlled via the amount ratio of acrylamide and bis-acrylamide as previously described^31^. Once both coverslips were ready, freshly made 10% ammonium persulfate (APS) and tetramethylethylenediamine (TEMED) were added to the polyacrylamide and bis-acrylamide mixture to initiate gel polymerization. Immediately after, a 2.5 μL drop of the mixture was put on the treated 22-mm square glass coverslip and topped with the FN patterned surface of the 12-mm round coverslip. The assembly was polymerized for 45 min before removing the round coverslip. The unpolymerized acrylamide was removed by rinsing twice with PBS. The resulting patterned gels were sterilized under 354 nm light for 5 min prior to adding the cells. The cells were seeded on top of the gels and allowed to adhere for 30 min. Unattached cells were washed off to avoid overgrowth of the patterns. The medium was reconstituted and the cells were incubated overnight until they reach confluency in the patterned regions.

### 2.2 Cell culture

Human vascular umbilical endothelial vein cells (HUVECs) purchased from Lonza (Basel, Switzerland) were cultured on M199 supplemented with 10% (v/v) endothelial growth medium (Cell application, San Diego, California), 10% (v/v) FBS (Lonza), 1% sodium pyruvate, 1% L-glutamine and 1% penicillin-streptomycin (Gibco, Waltham, MA). The cell plasma membranes were stained with CellMask (Thermo Fisher, Waltham, MA) to corroborate that the cells were forming an island of the desired size and shape.

### 2.3 Preparation of ICAM mAb-coated microspheres

Anti-ICAM coated microspheres were prepared as previously described^15^. After washing with PBS, 20-μm-diameter polystyrene microspheres (Polyscience) were treated overnight with glutaraldehyde for cross-linking while keeping them at 4°C. Subsequently, the microspheres were treated at room temperature for 6 hr with 200 μg/mL ICAM1 mAb (G-5, Santa Cruz Biotechnologies). Then, the microspheres were washed 3 times with PBS. The ICAM1 mAb-Coated beads were used immediately or stored at 4°C for several days before use.

### 2.4 Imaging

We acquired z-stacks of fluorescent images with 560-605 TRITC filter containing 40 planes with a separation of 0.2 μm between planes. In addition, we acquired a 650-684 Cy5 image to confirm that the cells were restricted to the patterned area. For the acquisition, we used a confocal scanning microscope (Olympus IX81, Shinjuku, Tokyo, Japan) with a cooled CCD (Hamamatsu, Hamamatsu, Shizuoka, Japan) using Metamorph software (Molecular devices, Sunnyvale, California) and a 40 X NA 1.35 oil immersion objective. After image acquisition, the cells were detached using trypsin (Acompany, Acity, Astate).

When imaging cell islands larger than the field of view of the microscope (i.e. 170um x 170um), we acquired several stacks to cover the whole micro patterned area. A ∼20% overlap in x and y was used between neighboring stacks to facilitate a posteriori merging. Overlapping stacks were combined using the “Grid/Collection stitching” plug-in of ImageJ (NIH).

### 2.5 Image Cross-Correlation for Deformation Measurement

We measured the deformation of the substrate in three dimensions by cross-correlating each fluorescence z-stack with a reference z-stack in which the substrate was not deformed, similar to del Alamo et al^19^. The fluorescence z-stack in the deformed state was compared with a reference z-stack which was recorded at the end of experiment, after the cells were detached and the elastic substrate recovered its undeformed state. The comparison between the deformed and undeformed (reference) conditions was performed by dividing both z-stacks into 3D interrogation boxes and maximizing the cross-correlation between each pair of interrogation boxes.

The accuracy of the image cross-correlation method used in del Alamo et al’s^19^ was refined here by introducing two improvements in their algorithm. First, we implemented the well-established multi-grid approach in which the correlation algorithm is run several times with interrogation boxes of progressively smaller sizes^32^. In this multi-grid approach, the results from each correlation pass are used to displace the interrogation box of the reference z-stack prior to image correlation. This method significantly increases the overlap between deformed and reference interrogation boxes, and allows for using smaller interrogation boxes without compromising the signal-to-noise ratio, which increases the spatial resolution.

Second, we adopted a novel variational approach to enforce the condition of global mechanical equilibrium of forces and moments. For a linearly elastic substrate, force equilibrium implies that the average of the measured deformation field must be zero in the x-y plane^33, 34^. This condition is typically enforced prior to recovering the traction stresses by subtracting the average value of the measured deformation, which can be non-zero due to image drift and other experimental noise sources. While this step effectively enforces global mechanical equilibrium, it modifies the measured deformation by the same value in all the interrogation boxes of each z-stack, regardless of local image quality in each box. A sounder approach consists of formulating the maximization of the cross-correlation function as a constrained optimization problem. At a given interrogation box with x-y coordinates given by indices *i* and *j*, the cost function of the problem is

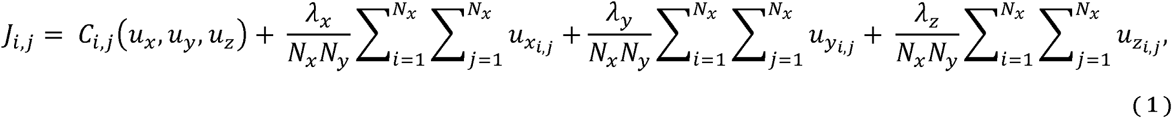

where *C*_*i,j*_ is the cross-correlation function between the deformed and undeformed interrogation boxes, *u*_*x*_, *u*_*y*_ and *u*_*z*_ are the displacement components in the x, y and z directions, and *λ*_*x*_, *λ*_*y*_ and *λ*_*z*_ are the Lagrange multipliers introduced to enforce that the spatial averages of *u*_*x*_, *u*_*y*_ and *u*_*z*_ are zero. To illustrate the method without loss of generality, we consider the one-dimensional case where

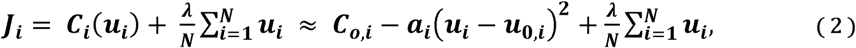

where the rightmost side of the equation includes a quadratic polynomial fit to the correlation function used for sub-pixel interpolation, and *u*_0,*i*_ is the deformation value that would be obtained for each box without enforcing global equilibrium. In this polynomial fit, the constant c_o,i_ represents the maximum value of *C*_*i*_ (*u*_*i*_), whereas *α*_*i*_ >0 indicates its convexity. If the signal-to-noise ratio of the image within an interrogation box is low, then *C*_*i*_ (*u*_*i*_) has a shallow peak and therefore *α*_*i*_ is small. Conversely, for interrogation boxes in regions of the image with high signal-to-noise, the correlation function has a sharp peak and the corresponding value of *α*_*i*_ is large.

Minimizing *J*_*i*_ with respect to *u*_*i*_ yields

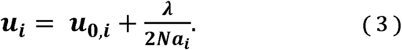

In order to find the value of the Lagrange multiplier, we enforce the global equilibrium condition 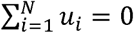, which results in 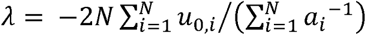. Plugging this result into equation 3 provides the solution to the constrained optimization problem,

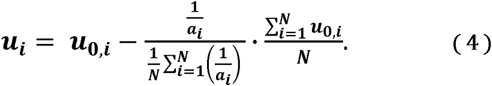

This result differs from the conventional method of removing the average, i.e. 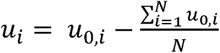, which applies the same correction to the displacement for all interrogation boxes. The variational approach proposed here applies a small correction in those interrogation boxes with higher signal-to-noise ratio, where 1/*α*_*i*_ is smaller than its average value across the whole image,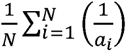. In the ideal case of an image of constant *α*_*i*_, the variational approach becomes equivalent to removing the average deformation. It is important to note that the variational approach can be embedded into the sub-pixel interpolation routine and does not require additional calculations of the cross-correlation function. Thus, it does not imply a significant increase in image processing cost.

Equilibrium of moments can be enforced in a similar way by including additional Lagrange multipliers in the cost function. To illustrate the procedure, we consider the moment around the y-axis caused by normal stresses *T*_*z*_ over the whole field *Ω*, which is

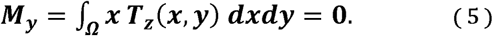

In Fourier space, this equation becomes^34^

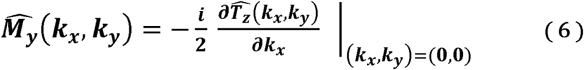

where 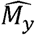 and 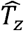 are the Fourier coefficients of *M*_*y*_ and *T*_*z*_, (*k*_*x*_, *k*_*y*_) is the wavenumber vector in the *x* and *y* directions, and *i* is the imaginary unit. The Fourier coefficient of the traction stresses is equal to the product between the coefficients of the measured deformation vector, *û*(*k*_*x*_, *k*_*y*_), and the tensorial Green’s function of the problem, *Ĝ* (*k*_*x*_, *k*_*y*_). The latter were given in closed analytical form by del Alamo et al^19^. Using the chain rule of derivation, the moment equilibrium constraint can be expressed as

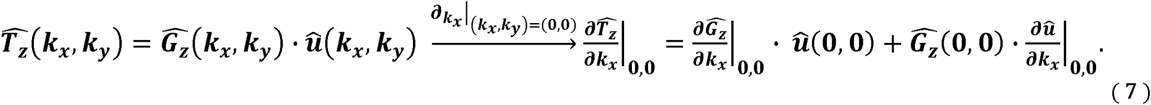

Force equilibrium implies that *û*(0,0) = 0, so the first term in the right-hand side of this equation vanishes. Bringing the remaining term back into physical space, we obtain the following expression for the spatial average of the y-moment,

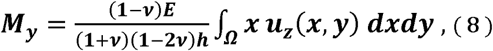

where *u*_*z*_ represents the normal deformation and 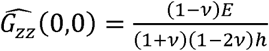 is the only non-zero term of the Green’s function for *T*_*z*_ ^19^. The above expression allows us to compute *M*_*y*_ directly from the measured deformation without having to solve for the traction force field. This is convenient in order to impose the constrain *M*_*y*_ =0 in our variational method via an additional constraint,

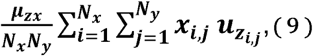

where *µ*_*zx*_ is the corresponding Lagrange multiplier. The same reasoning can be followed to calculate all the contributions to the x and y moments from all the components of the measured deformation vector. Adding Lagrange multipliers to enforce that all these contributions are zero leads to the following cost function:

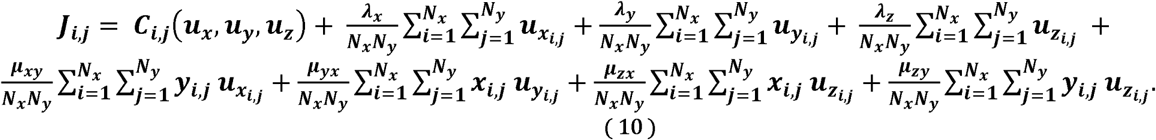

This cost function was minimized numerically using a gradient descent method in MATLAB (Mathworks, Natick, MA).

Figure 2 compares substrate deformations obtained by standard image correlation, multigrid image correlation, and variational image correlation with force and moment equilibrium constraints. Representative examples obtained for two different island sizes (50-and 90-micron diameter) are shown. Of note, the lateral deformations (*u, v*) obtained by standard correlation have discontinuities in some regions, which appear as “holes” or sharp changes in the sign of the deformation. These regions are indicated by arrows in Figure 2 and coincide with locations of large *u* and *v*, where a significant fraction of the fluorescent markers in each interrogation box move beyond the boundaries of the box as the substrate is deformed. Consequently, standard image correlation yields spurious results in these regions, whereas multigrid image correlation is able to better capture the large motion of the markers by progressively resizing and displacing the interrogation boxes. Additionally, both the standard and multigrid image correlation methods yield appreciable tilt in the normal deformation (*w*), which is evidenced by non-zero values of *w* outside and far from the edges of the cell islands and also in the profiles of normal deformation (w) along the cross-section denoted by the white dashed line. Figure 2 shows that these artifacts disappear if the correlation between the deformed and deformation-free reference image stacks is maximized subject to force and momentum equilibrium constraints.

**Figure 2.**
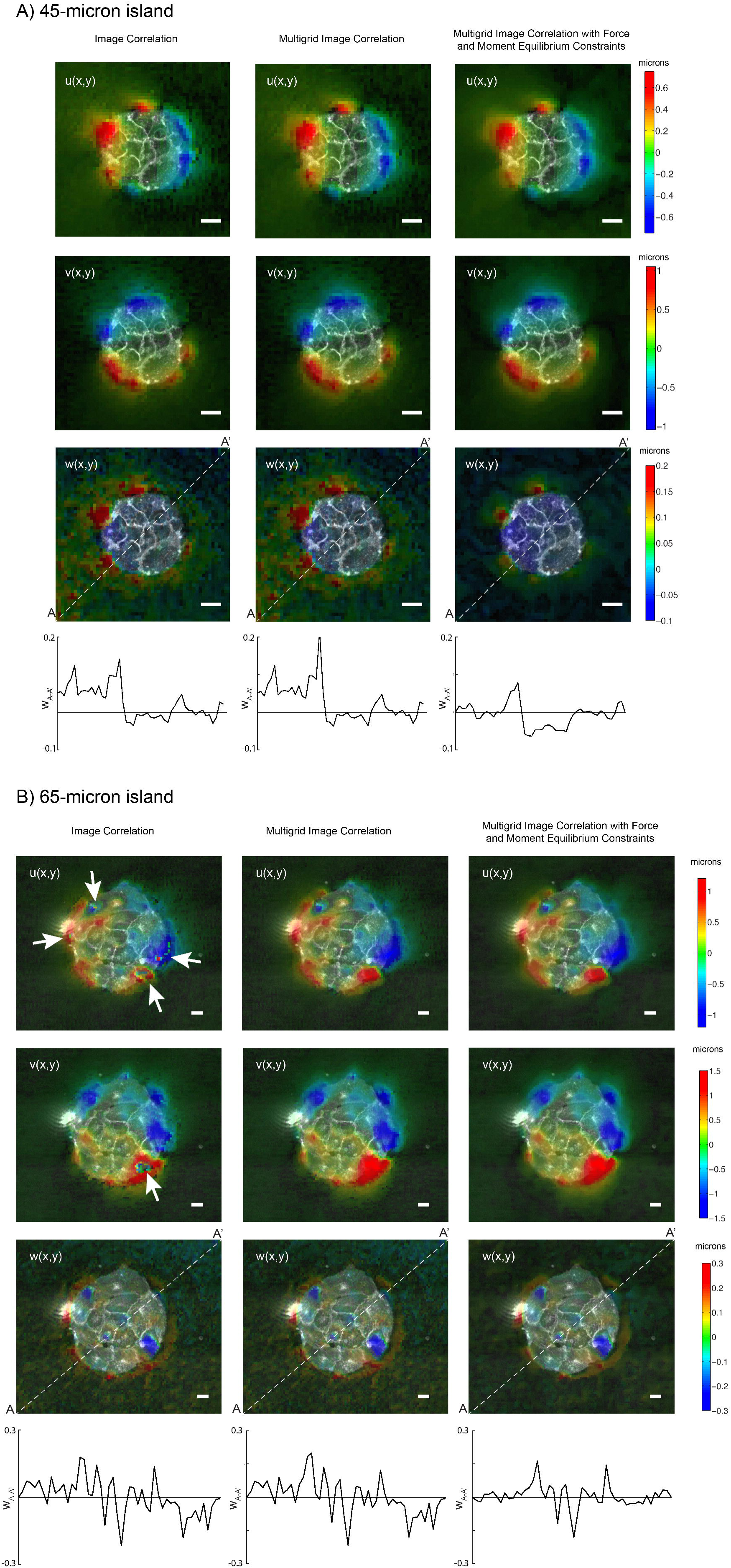
Illustration of the improvement gained by performing multigrid image correlation (center columns) and multigrid image correlation with force and momentum equilibrium constraints (right columns) compared to standard image correlation (left columns) for (A) a 45−μm radius island and (B) a 65−μm radius island. The measured displacement components in the x-direction (U), y-direction (V) and z-direction (W) are shown in the first, second and third rows respectively. The fourth row represents a side view of the measured W displacement along the line A-A’ (i.e. the white dashed line in third row).

The spatial resolution (i.e. the distance between adjacent measurement points) of the image cross-correlation algorithm is equal to 1/2 the size of the smallest interrogation window (i.e. the Nyquist spatial sampling frequency^32^). In the present experiments, we used interrogation windows of 32×32 pixels, yielding a spatial resolution of 2.7 μm. The sensitivity of the algorithm was enhanced by using standard sub-pixel interpolation of the image cross-correlation function^32^, yielding minimum detectable displacements of ∼0.02 μm.

### 2.6 Traction Force Microscopy

The 3D deformation of the PA substrate was measured at its top surface on which the cells were attached, following the method described above. Using these measurements as boundary conditions, we computed the three-dimensional deformation field in the whole polyacrylamide substrate by solving the elasto-static equation as previously described^19^. We then computed the traction stress vector exerted by the cells on the substrate, 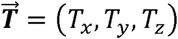. The spatial resolution of 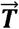 was 2.7 μm.

### 2.7 Equations of Mechanical Equilibrium in the Monolayer

#### 2.7.1 Lateral Deformation Problem

The computation of the intracellular stresses induced by lateral deformation follows the mathematical model formulated by Tambe *et al*^35^. The monolayer is assumed to be a thin, linearly elastic plate of uniform thickness *h* and Poisson ratio *v*. We use a coordinate system in which the x and y directions are parallel to the monolayer plane, while *z ∈* [−*h*/2, *h*/2] denotes the distance across the monolayer thickness (Figure 1A). Static equilibrium of forces in x and y yields

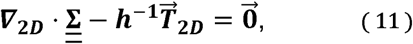

in the domain *Ω* occupied by the monolayer. In this equation, the tensor 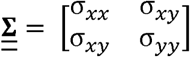 represents the intracellular stress induced by the lateral deformation of the monolayer, and the vector 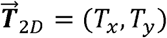 represents the tangential traction stresses at the interface between the cell monolayer and the substrate. The problem is closed via the Michel-Beltrami compatibility condition for continuity of deformation,

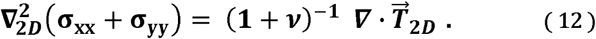

Researchers are often interested in computing just the intracellular tension (or compression, if it has negative sign) induced by in-plane monolayer deformation, *i.e.* the first invariant of 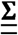 defined as 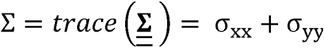, instead of the whole intracellular stress tensor^36, 37^. In that case, the Michel-Beltrami condition provides with a scalar Poisson equation to compute *S* from the traction stresses in an inexpensive manner.

Stress-free boundary conditions are prescribed at the edge of the cell aggregate (Figure 1E),

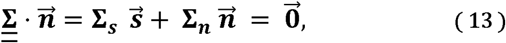

where 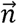 and 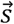 are respectively unit vectors normal and tangential to the edge of the monolayer, and Σ_*n*_ and Σ_*s*_ are the normal and tangential components of the stress vector. This boundary condition implies that Σ_*n*_ = Σ_*t*_ = 0.

#### 2.7.2 Bending Problem

In a uniform thin elastic plate subject to both lateral deformation and bending, the intracellular tension is given by

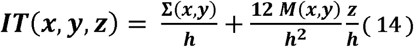

According to this expression, the intracellular tension caused by lateral deformation is constant across the shell thickness and proportional to Σ. In addition, bending has a contribution that varies linearly with *z* and is proportional to the moment function 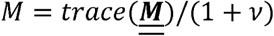, where 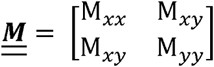 is the bending moment tensor^16^. To compare the magnitudes of the intracellular stresses caused by lateral deformation and by bending, it is useful to rewrite the above equation as

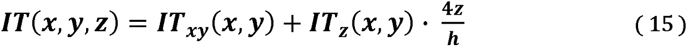

where *IT*_*xy*_ are the intracellular stresses caused by lateral deformation, and

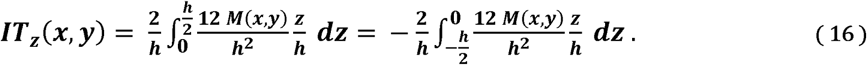

Thus, *IT*_*z*_ (*x, y*) = 3*M*(*x, y*)/*h*^2^ represents the average magnitude of the bending-induced intracellular stresses across the top and bottom halves of the shell. The calculation of *IT*_*z*_ is based on the equation of mechanical equilibrium of bending moments^16^,

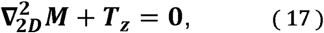

which relates the bending-induced intracellular tension to the normal traction stresses. The boundary conditions for this Poisson equation are obtained by imposing zero effective transverse shear stress (*V*_*n*_ = ∂_*n*_ *M*+ ∂_*s*_ *M*_*sn*_ = 0), and zero twisting moment (*M*_*sn*_ = 0) at the free edge of the monolayer^38^. These edge constraints are expressed mathematically as a homogeneous Neumann condition, ∂_*n*_ *M* = 0, since the zero-twist condition implies that the tangential derivative of *M*_*sn*_ is zero along the free edge of the monolayer.

For this Neumann boundary condition to be compatible with the Poisson equation (17), the cell monolayer must be in equilibrium of normal forces. Integrating (17) over the domain *Ω* occupied by the cells, and applying Gauss’ divergence theorem, we obtain

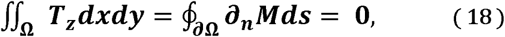

where *ds* is an element of line along the boundary of the cell domain, ∂*Ω*, and we have taken advantage of the fact that the closed-line integral ∮_∂Ω_ ∂_*s*_ *M*_*sn*_ *ds* is zero. This result implies that the bending problem is not solvable unless *T*_*z*_ has zero average in the cell domain, and underlines the relevance of variational image correlation methods that enforce global mechanical equilibrium of forces and moments (see section 2.4). Hereinafter, the intracellular tension caused by lateral deformation IT_xy_ will be referred to as “lateral tension”, and the intracellular tension due to bending deformation IT_z_ as “bending tension”.

### 2.8 Numerical implementation of boundary conditions at the edges of the monolayer

A level-set immersed interface method was used to enforce the boundary conditions at the edges of the monolayer^20^. This procedure allows for the numerical implementation of boundary conditions on arbitrary geometries while keeping a uniform Cartesian grid, which facilitates the discretization of the MSM equations by Fourier expansions. The Fourier discretization is computationally efficient and makes MSM interface easily with previous Fourier traction force microscopy methods^18, 19, 30, 34, 39, 40^.

In the immersed interface method, the geometry of the cell monolayer is defined implicitly through the introduction of a level set function, *Γ*(*x, y*), that takes different values the interior and exterior of the monolayer. In our calculations, we used a Heaviside function with *Γ* = 1 inside *Ω* and *Γ* = 0 outside *Ω* (Figure 1E) to augment the equation of tangential force balance as follows

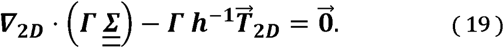

Considering that 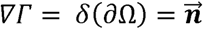 where *δ* represents a Dirac Delta and ∂Ω represents the boundary of the cell monolayer, the previous equation is equivalent to

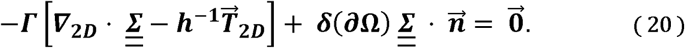

The first term in the left-hand side of this equation imposes tangential force balance in the region defined by *Γ*(*x, y*) *= 1, i.e.* inside the cell aggregate domain Ω. The second term imposes a stress-free boundary condition at the edge of cell aggregate, where *δ*(∂Ω) is non-zero. Note that because Ω and ∂Ω do not intersect, equation 20 independently imposes tangential force balance and the boundary conditions for the cell aggregate.

The bending problem can be treated in a similar manner, leading to the augmented equation

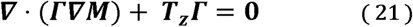

which is equivalent to

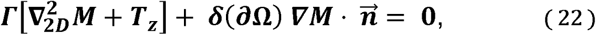

thereby recovering the equilibrium partial differential equation inside the cell aggregate and imposing the zero-shear-force condition, ∇Γ· ∇M = ∂_Il_ M= 0, at the edge of the aggregate.

### 2.9 Numerical Integration

The system of augmented equations of static equilibrium (19, 21) is linear but has spatially varying coefficients, which complicates solving it using a Fourier Galerkin approach (i.e. projecting the equations onto each Fourier mode to obtain an almost-diagonal linear system of equations). To overcome this difficulty, the system was solved iteratively using a dynamic relaxation technique^41^. In this approach, the equations are reformulated to represent the time-dependent dynamics of a vibrating shell, and are marched in time until the steady state corresponding to static equilibrium is reached. The dynamical relaxation equation for the lateral problem reads

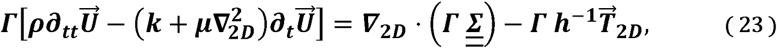

where 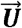 is the lateral deformation field of the cell monolayer, *ρ* is an arbitrary density, and a damping term with constants *k* and µ is introduced to accelerate convergence. Upon convergence, all time derivatives are zero, recovering the equations of static equilibrium and boundary conditions for the cell monolayer (19). Using the definition of strains as a function of the deformation and Hooke’s law, the time-dependent equation (23) can be recast as

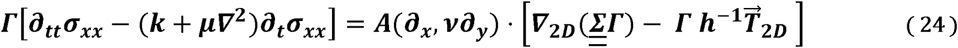

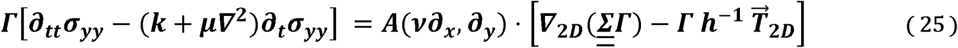

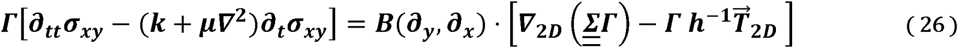

which only depend on the lateral stress tensor 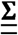, and where *A* = (1 − *v*^2^)^−1^ and *B* = −2(1 + *v*)^−1^. The bending problem is treated in a similar manner, which yields the relaxation equation

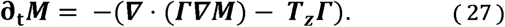

The minus sign in front of the right-hand-side of the equation does not affect the final solution upon convergence but it is necessary to keep the iteration numerically stable.

Equations (23-27) were discretized in a rectangular domain large enough so that the edges of the monolayer were separated at least 60 μm from edges of the image, using a Cartesian grid with inter node spacing Δ*x* = Δ*y* = 2.7 μm. The discretized dynamic relaxation equations were advanced in time using an explicit Euler integration scheme until the solution changed by less than 5% between consecutive iterations. As initial condition for the iteration, we used the solution obtained without immersed boundary forcing (i.e. *Γ* = 1 for the whole computational domain), which can be easily obtained by the Galerkin method. As is customary in level-set methods^42^, we used a smeared-out Heaviside indicator function instead of the sharp one in our numerical routines, in order to avoid spurious numerical oscillations in the results. Specifically, we convolved *Γ* with a Gaussian filter of width Δ equal to a few pixels wide. The precise value of Δ was chosen in each case to minimize the error of the solution based on our validation results, shown below.

## 3 Results

This section presents the results from validation analyses and experimental measurements of monolayer tension induced by 3D traction stresses in micro-patterned monolayer islands of varying shapes and sizes. Additionally, it illustrates how 3D-MSM can be applied to quantify the alterations in monolayer tension caused by localized perturbations in non-patterned monolayers.

### 3.1 Validation

We validated the numerical discretization and integration methods presented above for a plate under the synthetic tangential load

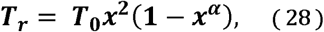

which resembles our experimental measurements of the traction stress under circular cell monolayer islands. In the above equation, *x = r/R* is the distance to the center of the island normalized by island radius (*R)*, and *T*_0_ is a characteristic stress. The parameter *α* ≥ 1 defines the width of the ring at edge of the cell island where the traction stresses are concentrated (see Figure 3A). This width can be quantified by the length scale 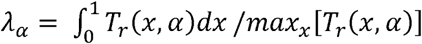, which is shown to decrease with *α* in the inset of Figure 3A.

**Figure 3.**
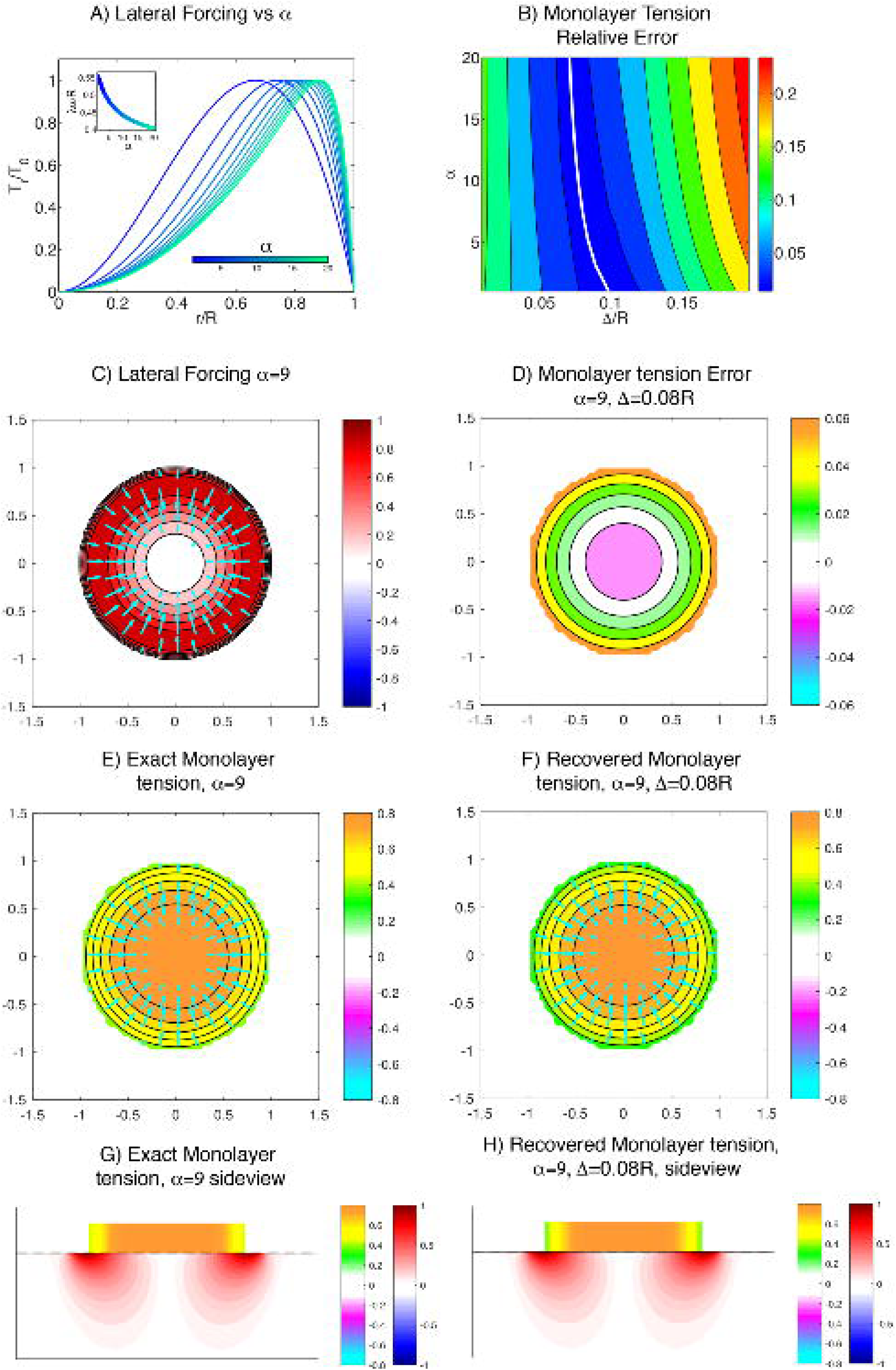
(A) Profiles of the tangential traction stress load used to validate 3DMSM for varying width of the ring where traction forces are concentrated. Inset: lengthscale of the width of the ring *λ*_*α*_ as a function of parameter *α*. (B) Average error 3DMSM in the simulation as a function of the smoothness of *Γ* (defined by the spatial filter width Δ/R), and the parameter that defines the width of the ring, *α*. (C) Example of a traction stress map for the particular case *α* = 9. (D) Error, (E) analytical and (F) numerical solution for the lateral intracellular stress. (G, H) Side view representation of the intracellular tension and substrate stress due to lateral deformation for both the analytical (G) and numerical (H) solutions. The black solid line represents the free surface of the deformed substrate. The red-blue colormap represents the magnitude 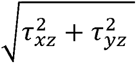 inside the gel. Orange-cyan colormap represents the intracellular tension.

In this axially symmetric case, the lateral elastic equilibrium and compatibility equations are reduced to

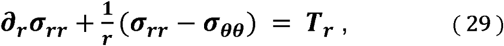

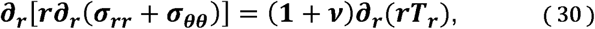

where (*σ*_*rr*_ and (*σ*_*θθ*_ are the diagonal elements of the lateral stress tensor expressed in polar coordinates. The exact analytical solution to these equations is

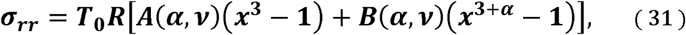

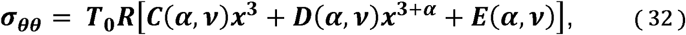

where the coefficients *A*(*α, v*), *B*(*α, v*), *C*(*α, v*) and *D*(*α, v*) are given in the Supporting Material, together with the derivation of the solution.

We determined the difference between the monolayer tension recovered from *T*_*r*_ using the procedures described in sections 2.6 – 2.8 above, and the exact monolayer tension obtained analytically. The difference is integrated over the cell island domain and normalized by the integral of the exact tension to yield the relative error:

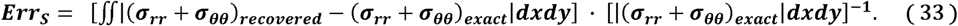

This error is plotted in Figure 3B as a function of the width of the low-pass filter used to smooth the Heaviside level-set function, *Δ*, and the parameter *α* that defines the sharpness of the input distribution of tangential traction stresses. The error analysis in Figure 3B indicates that the accuracy of the method is more sensitive to the filter width than to the spatial width of the traction stresses. The dependence of the error with *Δ* can be understood considering that *Δ* tends to zero the level set function becomes infinitely sharp, leading to spurious numerical oscillations in the results (Gibbs error)^42^. Conversely, if the filter is too wide it artificially smooths out the solution, which is manifested by an increase in the error. The optimal filter width is Δ ≈ 0.18*λ*_*α*_ (white line in Figure 3B), which corresponds to approximately 20% the spatial extent of the tangential traction stresses, and yields an error below 5%. This result allowed us to adjust the numerical routines in each specific experiment, since *λ*_*α*_ can be determined from the measured traction stresses.

Figure 3 also includes detailed results from a particular case corresponding to *α* = 9. The spatial distributions of the forcing traction stress and the absolute error in the monolayer tension are represented in Figures 3C and 3D, respectively. These data indicate that the error co-localizes with the forcing stresses. However, this error is small (<6%) and the spatial distributions of exact and numerically recovered monolayer tension are barely distinguishable (Figure 3E-F).

We used the same approach to validate our procedure to calculate the bending monolayer tension. We considered a circular cell monolayer island under the normal traction stress:

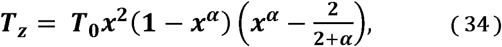

where the parameter *α* ≥ 1 again defines how concentrated the traction stresses are near the edge of the island. This synthetic traction stress distribution is shown in Figure 4A. It resembles the normal traction stresses measured in our experiments and satisfies global equilibrium of forces and moments, particularly of bending moments. The governing equation for this axially symmetric problem is

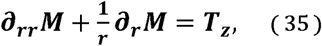

and the boundary condition is ∂_*r*_*M*|_*r*=*R*_ = 0. The exact analytical solution to this problem is

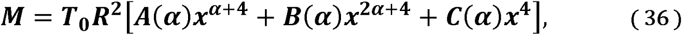

where the coefficients *A*(*α*), *B*(*α*) and *C*(*α*) are given in the Supporting Material, together with the derivation of the solution.

**Figure 4.**
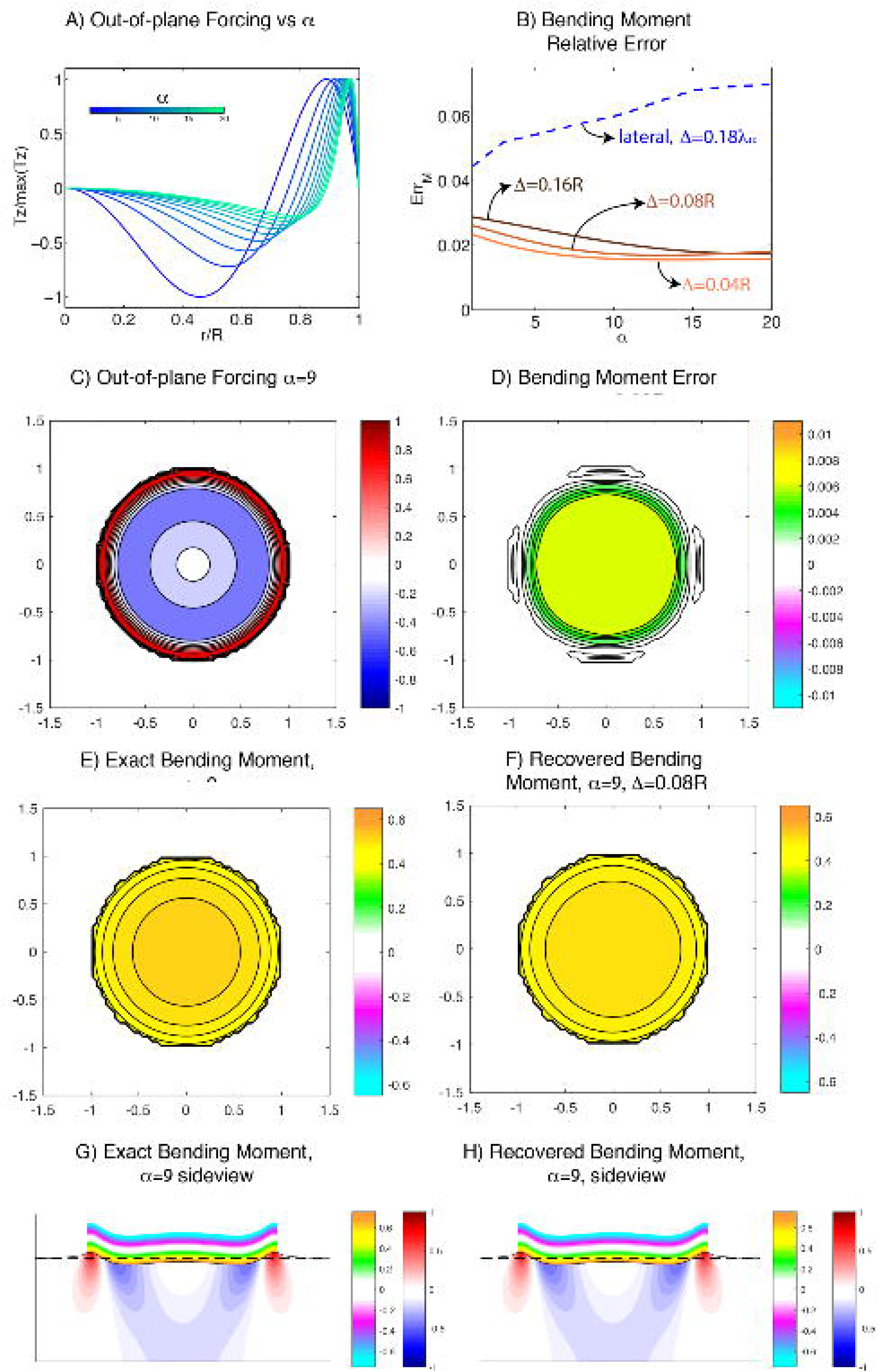
(A) Profiles of the normal traction stress load used to validate 3DMSM for varying width of the ring where traction forces are concentrated. (B) of the smoothness of *Γ* (defined by the spatial filter width Δ/R), and the parameter that defines the width of the ring, *α*. (C) Example of a traction stress map for the particular case *α* = 9. (D) Error, (E) analytical and (F) numerical solution for the bending intracellular stress. (G, H) Side view representation of the intracellular tension due to bending and *τ*_*zz*_ inside the gel for both the analytical (G) and numerical (H) solutions. The black solid line represents the free surface of the deformed substrate. Red-blue colormap represents inside the gel. The orange-cyan colormap represents the intracellular tension.

The relative error of the recovered bending moment is defined as

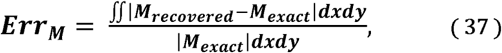

and is plotted in Figure 4B together with the lateral error obtained for the optimal value of the filter width, *Err*_*S*_ (*α*, Δ = 0.18*λ*_*α*_). Overall, *Err*_*M*_ is approximately independent of the shape of the forcing (α) and the width of the filter (Δ) used to smooth out the level set function Γ. Furthermore, *Err*_*M*_ is lower than *Err*_*S*_.

Similar to the lateral case, Figure 4 includes detailed results from a particular case corresponding to *α* = 9. Inspection of these data indicates that the magnitudes of both the bending moment and its absolute error are highest near the center of the island. As a result, the recovered bending moment (Figure 4F) underestimates the exact one near the center of the island, although this difference is small (relative error ∼2%).

### 3.2 Measurements of Lateral and Bending Monolayer Tension in Micropatterned Cell Islands

We used the methods described above to measure three-dimensional distributions of traction stress and intracellular stress in micropatterned islands of different shapes and sizes. Figure 5 shows a typical example of the measurements obtained for a circular island. The cells in the island collectively contract towards the island center by applying strong traction stresses in the x-y plane along the edge of the island (Figure 5A). In the normal direction (z), the cells pull upwards along the island edge, and the resulting force is balanced out by a weaker distribution of compressive stresses that is spread more uniformly over the whole extent of the island (Figure 5B). In cell islands with corners (e.g. triangular islands, see Figure 6A-B), both the contractile tangential traction stresses and the pulling z stresses are more concentrated near the high-curvature island corners, whereas the compressive stresses are still spread more uniformly over the whole extent of the island. These results are in general agreement with previously reported measurements of traction stresses in single-cell and collective-cell cultures^37,^ 43, 21.

**Figure 5.**
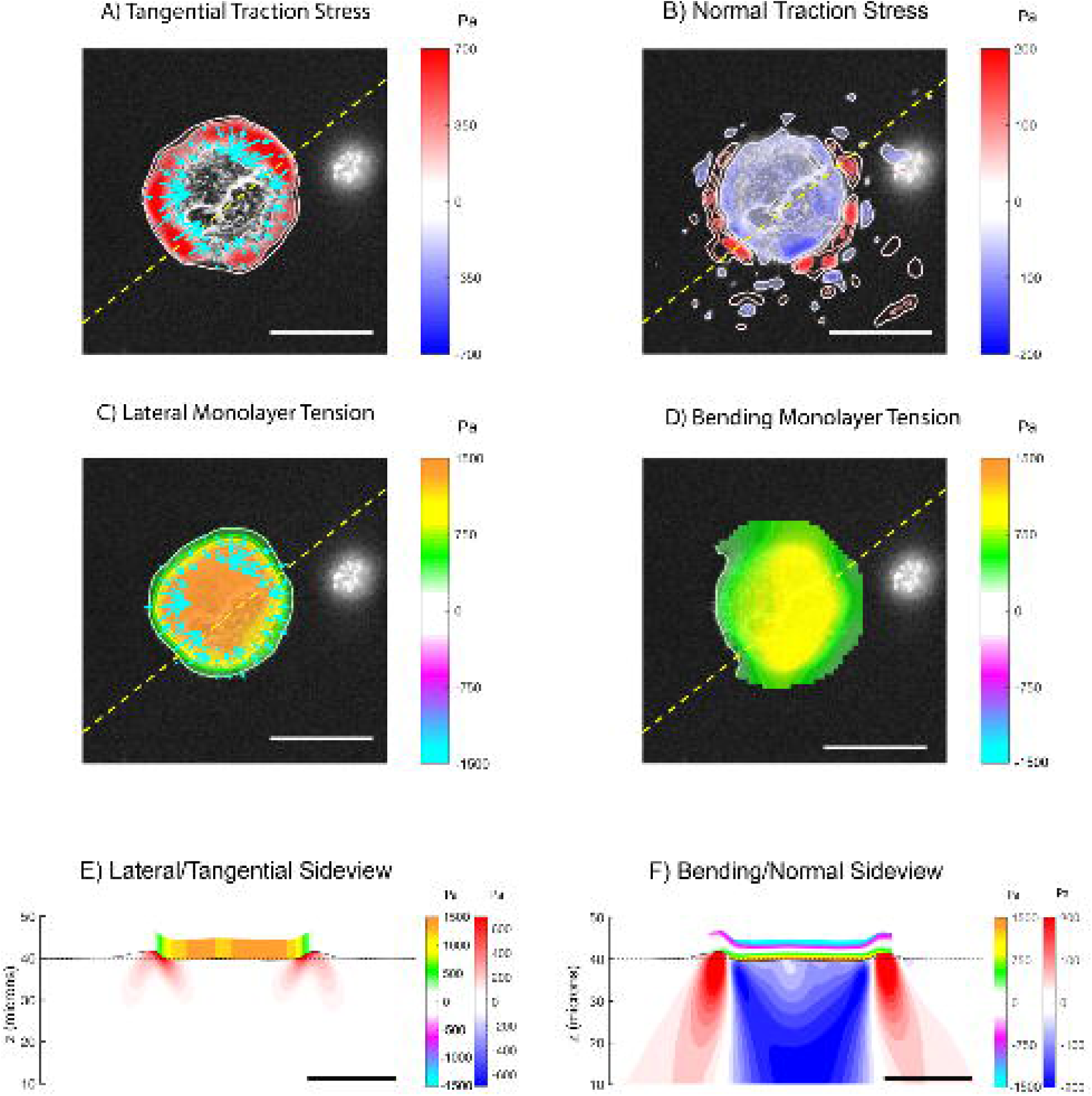
(A) Tangential traction stress (*τ*_*xz*_, *τ*_*yz*_) field on a 45-micron-radius cell island. The colormap represents magnitude of 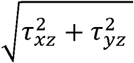. (B) Normal traction stress field, with positive values (red) pointing upwards and away from the substrate. (C) Intracellular tension due to lateral deformation (color), with tangential traction stress (arrows) superimposed for reference. (D) Intracellular tension due to bending stress. (E) Side view representation of the intracellular and substrate stresses caused by lateral deformation along a representative line (yellow-dashed lines in A, B, C, and D). The black solid line represents the free surface of the deformed substrate. The red-blue colormap represents the magnitude of 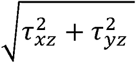 inside the gel. The orange-cyan colormap represents the intracellular tension. (F) Side view representing the intracellular tension due to bending stresses. Red-blue colormap represents the magnitude of *τ*_*zz*_ inside the gel. The orange-cyan colormap represents the intracellular tension due to bending. Scalebar = 50 μm.

**Figure 6.**
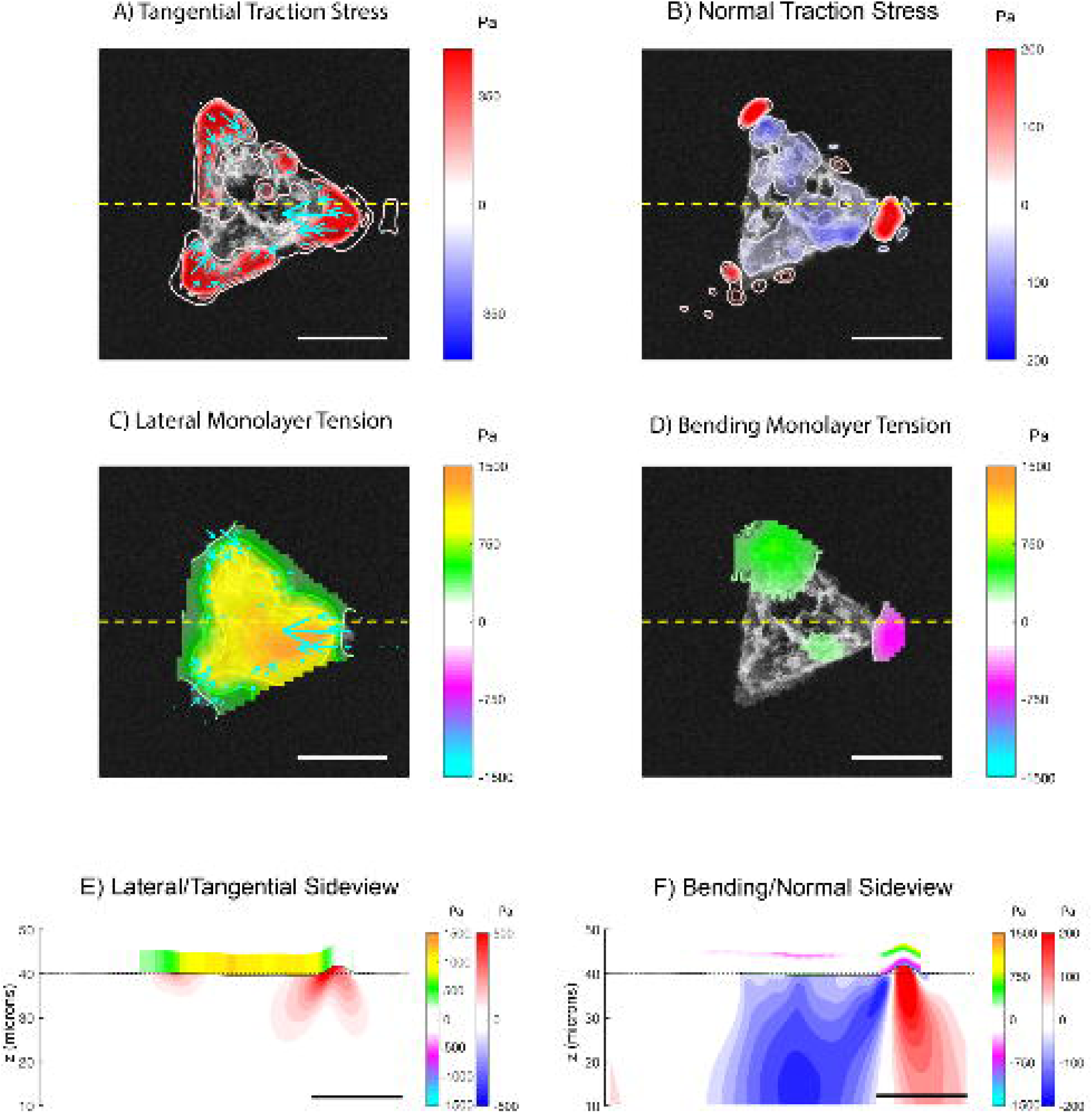
(A) Tangential traction stress (*τ*_*xz*_, *τ*_*yz*_) field on a triangular cell island of side 100 microns. The colormap represents the magnitude 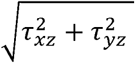. (B) Normal traction stress field, with positive values (red) pointing upwards and away from the substrate. (C) Intracellular tension due to lateral deformation (color), with tangential traction stress (arrows) superimposed for reference. The yellow dashed line indicates the section on which the side views are represented in panels E and F. (D) Intracellular tension due to bending. (E) Side view representation of the intracellular and substrate stresses caused by lateral deformation (corresponding to the yellow-dashed lines in panels A-D). The black solid line represents the free surface of the deformed substrate. The red-blue colormap represents 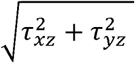 inside the gel. The orange-cyan colormap represents the intracellular tension due to lateral deformation. (F) Side view representing the intracellular tension due to bending. TherRed-blue colormap represents *τ*_*zz*_ inside the gel. The orange-cyan colormap represents the intracellular tension due to bending. Scalebar = 50 μm.

The fact that tangential and normal traction stresses tend to co-localize in our measurements suggests that there is a relationship between them. To test this hypothesis, we sampled the peak values of the tangential and normal traction stresses along 8 radial lines of each island (evenly spaced angularly by 45 degrees). The two variables are plotted in Figure S3M-A in the Supplementary Materials. The p-value of the null hypothesis that there is no correlation between them was p=3×10^−18^, their correlation coefficient was R = 0.67 and the lower and upper 95% confidence intervals of this coefficient were 0.56 and 0.75 respectively. Thus, the tangential and normal traction stresses are significantly correlated. Then, we determined the angle at which the focal adhesions pull vertically by calculating the arctangent of the ratio between the peak normal and peak tangential traction stress. The results, plotted in Figure SM3B in the Supplementary Materials, suggest that this angle is between 8 and 10 degrees. There is no statistical difference in pulling angles between the three island sizes.

The traction stresses were used as inputs to calculate the internal monolayer stresses caused by lateral deformation and the bending of the cell islands. The monolayer tension caused by each of these two contributions is also plotted in Figure 5 for an example circular island. It is important to note that the tensions induced by lateral deformation (Figure 5C, E) and bending (Figure 5D,F) have comparable magnitudes, especially in the region near the edge of the island. The tension coming from lateral deformation starts at low values at the free edge of the island and continuously increases towards the island center. This increase is sharpest near the edge of the island where the tangential traction stresses are maximum. The bending tension also rises sharply from its value at the island edge but, in contrast to the lateral tension, it plateaus in the interior region of the island. This behavior can be understood by considering that monolayer bending was highest near the island edge due to the concentration of positive and negative normal traction stresses in that region (Figure 5F).

The distribution of lateral tension was overall similar in triangular cell islands (Figure 6C,E), although we observed a trend for triangular islands to develop more pronounced asymmetries than circular islands (compare Figures SM3 and SM4 in the Supplementary Materials). This difference could be explained by considering that the stress distribution in triangle islands is more complex than in circles (compare Figure 3 with Figure SM4 in the Supplementary Materials). Thus, the cells in triangular islands were subjected to more heterogeneous mechanical environment which might promote the observed asymmetries.

#### 3.2.1 The effect of monolayer island size on the lateral and bending monolayer tensions

To study how cell island size affects the lateral and the bending monolayer stresses, we performed experiments on circular micropatterned islands of different radii (*R*_*island*_ = 25, 45, and 65μm). The choice of circular islands facilitates compiling statistics to quantify in detail the spatial distribution of the monolayer stresses. For each island, we expressed traction and monolayer stresses in a polar coordinate system with origin at the center of the island (e.g. the tension was denoted as *s*_*i*_ (*r, θ*) where the index *i* identifies islands within each diameter group, and *0<r<R*_*island*_ and 0<*θ* < 2π are the radial and azimuthal coordinates), and calculated their mean and standard deviation along the angular direction,

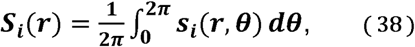

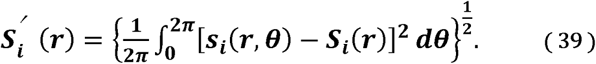

These variables were averaged for the total number of islands (N = 8, 4, and 5 for the 25-, 45-and 65-μm-radius islands), within each island radius group to obtain the mean stress profiles 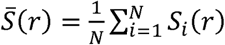, and the standard deviation profiles 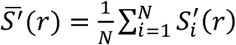. They are plotted in Figures 7 and 8 as a function of the distance to the edge, *d*_*edge*_ = *R*_*island*_ – *r*.

**Figure 7.**
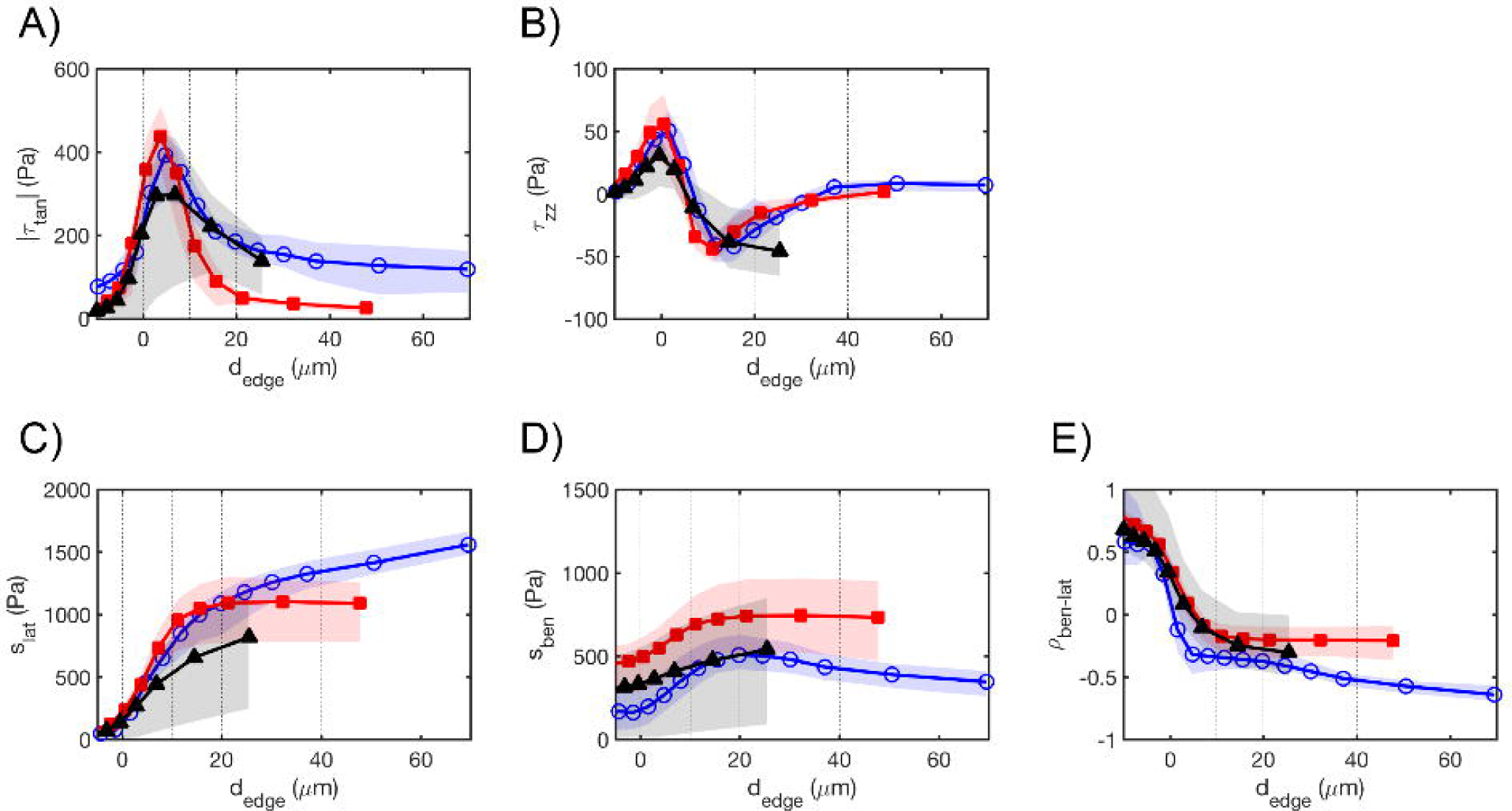
Mean profiles of: magnitude of tangential traction stress (A), normal traction stress (B), intracellular tension due to lateral deformation (C) and bending (D), and relative dominance of bending vs lateral intracellular tension (E); with respect to the distance to the edge in circular islands of 25(black)-, 45(red)-and 65(blue)-μm radius. The solid lines represent the mean values for each island size and the shaded regions represent the 95% confidence interval of the mean, computed using the bootstrap method with 1,000 iterations.

**Figure 8.**
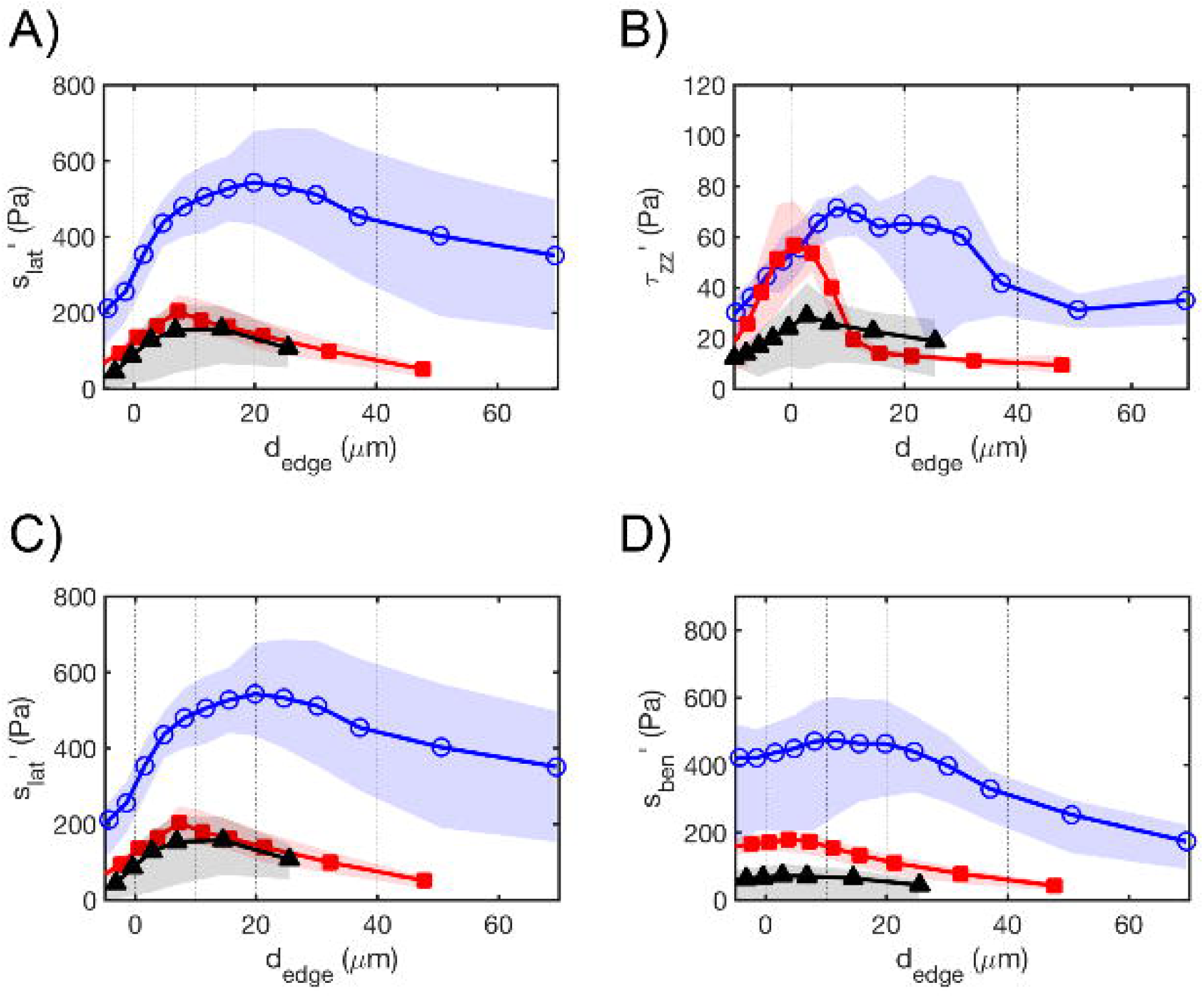
Profiles representing the azimuthal standard deviation (eq. 39) of: magnitude of tangential (A), normal (B) traction stress, and intracellular tension due to lateral deformation (C) and bending (D); with respect to the distance to the edge in circular islands of 25(black)-, 45(red)-and 65(blue)-μm radius. The solid lines represent the mean for each island size and the shaded regions represent the 95% confidence interval of the mean, computed using the bootstrap method with 1,000 iterations.

The mean profiles of tangential and normal traction stresses do not vary significantly with island size and only depend on the distance to island edge (see Table 1 in Supplementary Materials). The mean profiles of the tangential traction stresses (|*τ*_*tan*_| (*d*_*edge*_)) peak around ≈ 5 μm away from the island edge (Figure 7A). This behavior is independent of island size, suggesting that these traction stress patterns are mainly generated by the outermost single row of cells of the island. Consistent with previous observations on single cells^43^, the normal stresses (|*τ*_*zz*_| *d*_*edge*_)) have a maximum (i.e. pulling) that precedes the peak of the tangential stresses by ≈ 5 μm, and a minimum (i.e. pushing) that follows it by ≈ 10 μm. Legant *et al* {Legant, 2013 #82} suggested that the structural reason for this pattern could be that the focal adhesions are stiffer than their surroundings. According to their model, the adhesions would behave as a “rigid tile” that rotates when pulled by the cytoskeleton. This rotation would cause the part of the adhesion that is closer to the cell edge to displace upward, and the part of the adhesion that is closer to the cell interior to displace downward. Our results are consistent with that model.

The value of this maximum, *τ*_*zz,max*_ ≈ 60 Pa, is lower than the maximum value of tangential stresses |*τ*_*tan*_|_*max*_ ≈ 400 Pa for all island sizes, and this difference is statistically significant (p-values 0.001, 0.03 and 0.008 for the 25-, 45-and 65-μm-radius islands respectively). These peak values correspond with an upward pulling angle *γ* = tan^-l^(*τ*_*zz,max*_ /|*τ*_*tan*_|_*max*_) = 8.5 degrees at focal adhesions, which is in good agreement with the pulling angle statistics presented above. Away from the edge, the mean profiles of the traction stresses decay towards the center of the island, and this behavior is more evident for the larger islands. Apart from that difference, the mean profiles of tangential and normal traction stresses do not vary significantly with island size (see Table 1 in Supplementary Materials). Of note, the *τ*_*zz*_ profile remains negative-valued beyond its minimum, suggesting that in average the cells away from the edge exert a gentle pressure on their substrate (consistent with the data shown in Figures 5B, 6B and SI2B).

The mean profiles of lateral and bending tension, *s*_*lat*_ (*d*_*edge*_) and *s*_*ben*_(*d*_*edge*_), follow similar trends with *d*_*edge*_ regardless of island size (Figure 7C,D, Table 1 Supplementary Material). They raise sharply near the edge of the island where the traction stresses are strongest, and reach a plateau in the interior of the island. This behavior, which is more apparent for the larger islands, suggests that a monolayer reaches homeostasis a few cell rows far away from its free edge, not only for its lateral tension as previously reported^44^, but also for its bending tension. Remarkably, the lateral tension is zero at the edge of the islands (as dictated by the stress-free boundary condition) whereas the bending tension is not, which implies that there is a region near the island edge where bending is the dominant source of mechanical tension inside the monolayer. To quantify the relative contributions of lateral deformation versus bending to monolayer tension, we computed the quantity

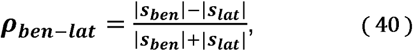

which varies between −1 (pure lateral distortion) and 1 (pure bending). The profile of *R*_*ben-lat*_ (Figure 7E) shows that bending is the dominant source of intracellular tension near the island edge. Towards the interior of the island, the lateral deformation becomes increasingly more important, but bending is still an appreciable source of intracellular stress at the center of the island (*ρ*_*ben-lat*_ ≈ −0.5). This behavior seems to depend little on cell island size, although we observe a trend for bending to become less important far from the edge of the island with increasing *R*_*island*_. (Table 1 Supplementary Material).

The standard deviation profiles in Figure 8 represent the fluctuations of the stresses with respect to the azimuthally-averaged profile of each island. Thus, they quantify the spatial variations in stress rather than island-to-island variability. These fluctuations generally increase with island size, both for the traction stresses and the monolayer stresses and both for lateral and bending deformations. The presence of larger stress fluctuations in larger islands can be noticed by comparing figures SM1 and SM2 in the Supplementary Materials.

### 3.3 Application of 3D MSM beyond cell islands

In cell islands, both tangential and normal traction stresses localize near the free edge of the cell monolayer, so that the most significant changes in intracellular tension occur near the monolayer edge. In order to illustrate the application of 3DMSM in a condition where the main source of monolayer bending is not located at a free monolayer edge, we seeded 20-μm-diameter microspheres on top of a non-patterned confluent monolayer of HUVECs. The microspheres were functionalized by coating them with anti-intercellular adhesion molecule-1 (ICAM-1). The expression ICAM-1 on the surface of VECs is known to trigger endothelial adhesion via integrins and other transmembrane receptors {Muller, 2014 #1; Nourshargh, 2014 #4; Vestweber, 2015 #5}. The microspheres were deposited far away from the edges of the culture dish to prevent edge effects. Recent work has shown that upon deposition, the functionalized microspheres are recognized by the HUVECs and firmly adhere to the apical surface of the monolayer {Yeh, 2018 #121}. Concurrently, the cell monolayer undergoes notable changes in tangential and normal traction stresses around the microsphere, as shown in the example case of Figure 9A-B. However, the effect of these changes in both lateral and bending monolayer tension was so far unknown.

**Figure 9.**
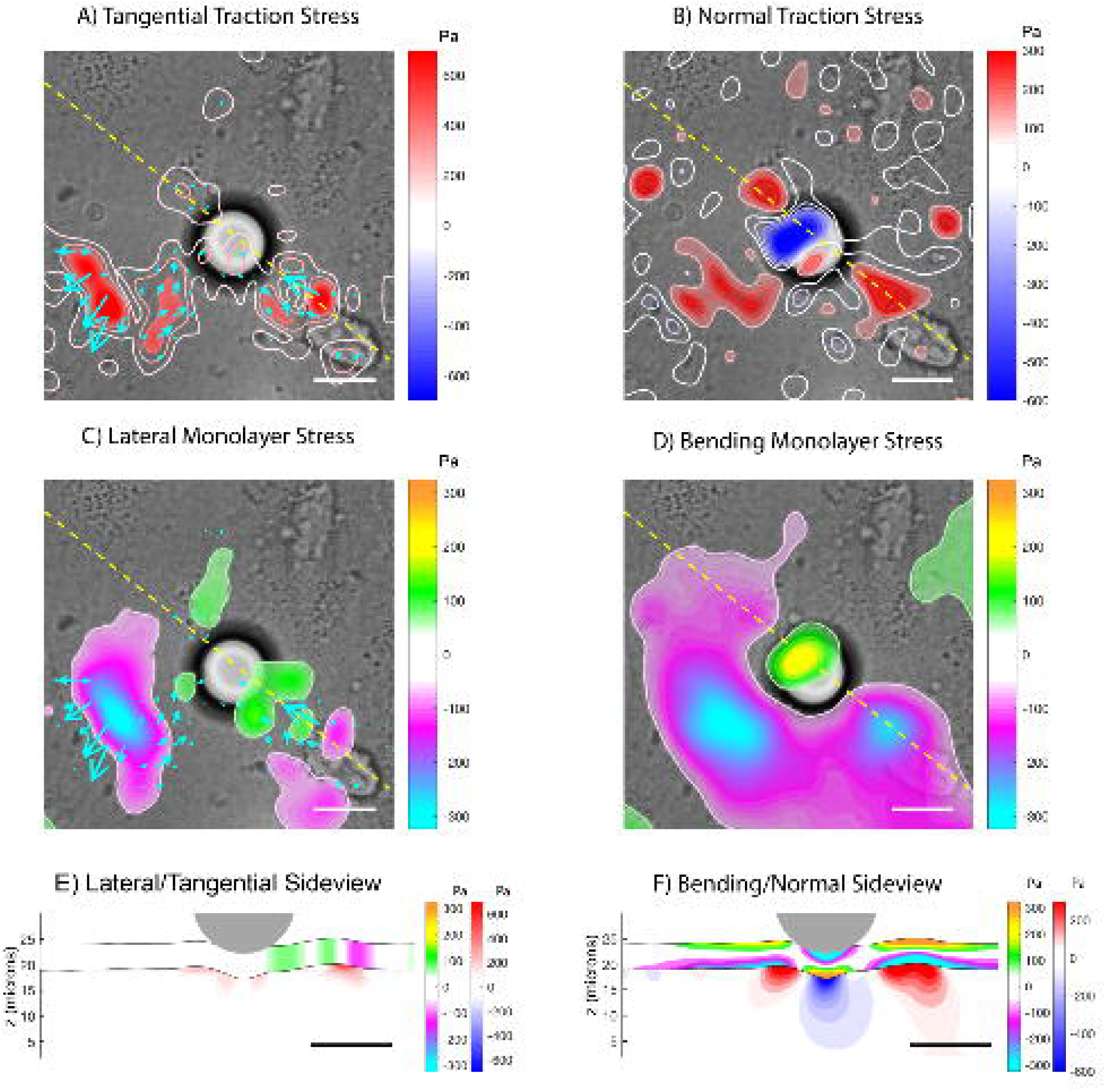
(A) Tangential traction stress (*τ*_*xz*_, *τ*_*yz*_) field of a cell monolayer under the presence of an ICAM-coated microsphere of 20 μm diameter. The colormap represents the magnitude 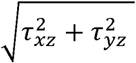. (B) Normal traction stress field, with positive values (red) pointing upwards and away from the substrate. (C) Intracellular tension due to lateral deformation (color), with tangential traction stress (arrows) superimposed for reference. The yellow dashed line represents the section on which the side views are represented in panels E and F. (D) Intracellular tension due to bending. (E) Side view representation of the intracellular and substrate stress due to lateral deformation (corresponding to the yellow-dashed lines in panels A-D). The black solid line represents the free surface of the deformed substrate. The gray circle represents the bead located on top of the cell monolayer. The red-blue colormap represents 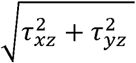 inside the gel. The orange-cyan colormap represents the intracellular tension due to lateral deformation. (F) Side view representing the intracellular tension due to bending. The red-blue colormap represents *τ*_*zz*_ inside the gel. The orange-cyan colormap represents the intracellular tension due to bending. Scalebar = 20 μm.

Using 3DMSM, we determined the changes in lateral and bending monolayer tension elicited by the microspheres. Figures 9C-F show that there were notable tension changes in the periphery of the microsphere. Notably, the bending tension increased forming a spatial pattern that co-localized tightly with the microsphere (Figure 9D,F). These changes in monolayer tension could locally modulate endothelial permeability.

## 4 Discussion

Monolayer Stress Microscopy (MSM) is becoming an increasingly widespread method to quantify the collective generation and transmission of intracellular stress. There are several different approaches to calculate intracellular stresses from measurements of substrate deformation, including the discrete application of Newton’s third law at cell-cell boundaries^45^, continuum mechanics models based on the analogy between the monolayer and a thin-plate^8, 35^, dissipative particle dynamics simulations^10^ and Bayesian inference^12^. However, none of these approaches considers that a cell monolayer can undergo bending due to mechanical stresses pointing in the direction perpendicular its surface, even when cultured on a flat substrate. These bending stresses can be generated by the cells that form the monolayer^9, 17^ but also by cells outside of the monolayer. For example, leukocytes undergoing endothelial transmigration generate mechanical forces that bend the endothelium at the transmigration site^15^. Furthermore, epithelial bending and invagination driven by mechanical forces play a key role in the development of most tissues and organs^14^. Currently, there is a lack of experimental data about the contribution of bending to intracellular tension, and also about the spatial organization of this mechanical signal throughout monolayers.

The intracellular tension caused by small bending in a cell monolayer can be inferred by extending Trepat et al’s^8, 35^ thin-plate analogy to consider not only equilibrium of forces, but also equilibrium of bending moments. The inputs to the extended 3-D MSM method are the 3-D traction stresses at the interface between the cell monolayer and its substratum, i.e. [*τ*_*xz*_ (*x, y*), *τ*_*xz*_ (*x, y*), *τ*_*zz*_ (*x, y*)], which requires interrogating the 3-D deformation of the substrate in a thin volumetric slice near the surface of the substrate^19^. Similar to traction force microscopy, where global force balance yields an integral constraint for the measured deformations (i.e. zero average displacement)^33^, global balance of bending moments leads to an additional constraint (i.e. zero average tilt). Given that the image processing algorithm for measuring substrate deformation involves the optimization of image cross-correlation, the constraints can be incorporated during image processing instead of during stress recovery. This approach is advantageous since it adapts the weight of the constraints in each region of the image according to its local signal to noise ratio, and it generally yields more realistic displacement measurements.

The MSM equations were discretized numerically on a Cartesian grid using Fourier series. The boundary conditions at the edges of the monolayer were imposed via a level-set immersed interface method^20^. In this formulation, the problem is solved in a rectangular domain that encompasses the edges of the monolayer, i.e. the immersed interface, and virtual body forces are imposed to enforce boundary conditions. This procedure leads to a relaxation problem that can be marched in time until it converges to a steady state solution that simultaneously satisfies equilibrium of moments and forces, as well as the boundary conditions at the edges of the monolayer. Compared to finite element methods, the Fourier immersed interface method does not require solving large linear systems of equations. Additionally, it interfaces exactly with the Fourier method used to determine the traction stress inputs to the MSM equations^18, 19, 30, 34, 39, 40^. The numerical solver was validated by prescribing a family of synthetic traction stress distributions for which the lateral and bending monolayer stresses can be calculated analytically, and which mimic our 3D experimental data on endothelial monolayer islands. This validation study suggested that the main source of numerical error is the level-set function used to impose the immersed interface boundary condition. When the level-set function varies too sharply, we observe spurious oscillations near the monolayer edge due to Gibbs error. On the other hand, if the level-set function is too smooth, the virtual body forces propagate inside the monolayer and the boundary conditions are smeared out. These two effects can be balanced out when the steepness of the level-set function is adjusted to be proportional to the steepness of the tangential traction stresses near the island edge, in which case the error decreases to ∼5%.

The thin-plate model assumes that the cell monolayer is a continuum of uniform thickness and linear material properties. To which extent these assumptions depart from reality, and what is their influence in the output of MSM are questions that deserve attention. The assumption of continuum medium is supported the reported agreement between MSM methods that model a cell monolayer as interacting particles and MSM methods that use continuum models, at least for lateral distortions^10^. It is reasonable to think the hypothesis of continuum should break down when the number of cells in the monolayer is sufficiently small (e.g. in small islands), and this is an area that deserves further investigation. Non-uniformities in material properties or monolayer thickness lead to concentration of stresses, an effect that is neglected in the model. Specifically, stresses concentrate in regions of locally lower thickness or higher Young’s modulus (*E*) and this effect is inherently the same for lateral and bending stresses. In principle, both effects could be incorporated into the model if one knew the spatial distribution of *E* and *h* across the monolayer, but these quantities are difficult to measure reliably. Available data suggests that cell monolayers have approximately uniform thickness ^46^, and the effects non-uniform material properties have been previously shown to be modest in two dimensions ^35^. Emerging MSM techniques are focused on bypassing the limitations of the thin plate model by developing methods to infer not monolayer stresses but also its material properties ^47-49^. These approaches shall be incorporated into three-dimensional MSM in the future.

Within the framework of the thin plate model, the intracellular tension induced by convex bending (*M>0*) varies across the thickness of the monolayer, reaching its maximum (tension) and minimum (compression) values at the top (i.e. apical) and bottom (i.e. basal) surfaces. This variation is given by

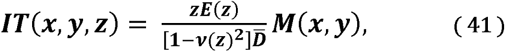

where 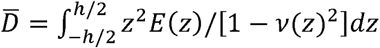 is the flexural rigidity of the monolayer averaged across its thickness. The opposite distribution of bending stresses (i.e. apical compression and basal tension) is obtained for concave bending (*M<0*). The z-dependence of the bending-induced intracellular stresses makes it difficult to quantify their average magnitude across monolayer width. In the idealized case of uniform material properties, the value of the bending stresses averaged across monolayer width is zero, so we chose the mean of their absolute value to quantify their magnitude. We note, however, that the mean bending stresses across the thickness of the monolayer need not be zero in the more realistic scenario where the mechanical properties of cells vary in the *z* direction, due to stress concentration. Recent data indicate that the main tension bearing elements of adherent cells, i.e. the stress fibers, have different orientations in the basal and apical regions of cells adhering to convex surfaces^50^. The reported orientation patterns are consistent with the apical stress fibers but not the basal ones bearing tension, in agreement with bending creating mechanical tension in the apical surface and compression in the basal surface. How cells are able to withstand compressive stresses is less clear, although the nucleus and cytoplasmic pressure mediated by membrane curvature have been postulated as candidates^50, 51^. The fact that bending may result in compressive or tensile stress and deformation in different compartments of the cell based on their apical/basal localization could have implications for mechanotransduction.

In epithelial cell sheets, the intracellular stresses caused by lateral cell contractility have been shown to extend over distances much longer than the size of a single cell, and to contribute to collective mechanosensing^21, 52^. However, the spatial organization of intracellular stresses caused by normal contractility has not been explored systematically so far. To address this question, we applied 3D-MSM to confluent micropatterned cell islands of different shapes and sizes, and obtained the first measurements of intracellular tension caused by out-of-pane traction stresses. The use of micropatterned islands allowed us to the prescribe boundary conditions at the free edge of the monolayer based on first principles. Micropatterning extracellular matrix proteins onto the substratum provides a robust way to culture cell islands within a region of tunable size and shape^53^. Furthermore, micropatterning allows for repeatable geometric conditions, thus facilitating the statistical analysis of the experimental data.

Our results indicate that cellular traction forces concentrate at the edges of monolayer islands. In circular islands, lateral inward pulling stresses are quasi-uniformly distributed along the island perimeter thus balancing out to zero net force. In the normal direction, force equilibrium is achieved by a ring of upward pulling stress at the island edge surrounding a downward pushing ring. These data are consistent with previous models where collective cell cooperativity at the mechanical level is achieved by tight transmission of stress across cellular junctions in the interior of a monolayer, together with cell-substrate traction stresses concentrated at the monolayer edges ^17, 21^. While the observed stress patterns were overall reproducible, we found that stress fluctuations increase with island size. It seems reasonable to expect this behavior because the configuration entropy of the monolayer increases as more cells fit inside larger islands. Previous models ^17^ have implied that heterogeneities in cell-cell junctions may lead to higher traction stress fluctuations inside the monolayer islands. Our measurements appear to be consistent with this idea.

Our measurements revealed that the magnitude of bending intracellular tension can exceed that of the tension caused by lateral contractility near the edges of a cell monolayer (particularly in regions of high edge curvature like corners), or in interior regions where the monolayer is subjected to biomechanical perturbations. We also found that the internal stresses caused by bending remain spatially confined in comparison to the internal stresses caused by lateral contractility, and tend to balance out within a few (∼1) cell lengths into the monolayer. This confinement might be relevant for phenomena like cellular extravasation, which are accompanied by significant normal deflections of the endothelial monolayer, and which require spatially regulated changes in endothelial permeability^15 54^. Our experiments with functionalized microspheres indicate that biomechanical perturbations to endothelial monolayers far away from the monolayer edge cause significant local changes in bending tension, and illustrate the application of 3D MSM to study biological processes.

## AUTHOR CONTRIBUTIONS

R.S., A.A., J.L., S.V., and J.C.d.A. designed the study. A. A. and Y.-T. Y. performed the experiments. R.S. and J.C.d.A. developed the method and analyzed the data. R.S., A.A., J.L., S.V., and J.C.d.A. wrote the manuscript.

