## supplementary text and figures for "Three-dimensional Monolayer Stress Microscopy"

### SUPPORTING MATERIAL

#### Derivation of analytical solutions for validation

Lateral Problem: We validated our procedure to recover monolayer tension from tangential tractions stresses for a cell monolayer island under the radially symmetric traction stress:

$$T_r = T_0 x^2 (1 - x^\alpha), \quad \text{(A1)}$$

where *r* is the distance to the center of the island, *R* is the island radius and $x = r/R$. The parameter $\alpha \geq 1$ defines how concentrated the traction stresses are near the edge of the island, and $T_0$ is a characteristic stress. This synthetic traction stress distribution is in equilibrium of forces and moments. The governing equations for this problem are the lateral elastic equilibrium and compatibility equations, i.e.

$$\partial_r \sigma_{rr} + \frac{1}{r}(\sigma_{rr} - \sigma_{\theta\theta}) = T_r, \quad \text{(A2)}$$

$$\partial_r[r\partial_r(\sigma_{rr} + \sigma_{\theta\theta})] = (1 + \nu)\partial_r(rT_r), \quad \text{(A3)}$$

where $\sigma_{rr}$ and $\sigma_{\theta\theta}$ are the diagonal elements of the lateral stress tensor expressed in polar coordinates. The boundary condition for this problem is zero radial intracellular stress at the island edge $\sigma_{rr}(x = 1) = 0$. Given the form of these equations and the boundary condition, we try solutions of the form

$$\sigma_{rr} = T_0 R[A(\alpha, \nu)(x^3 - 1) + B(\alpha, \nu)(x^{3+\alpha} - 1)], \quad \text{(A4)}$$

$$\sigma_{\theta\theta} = T_0 R[C(\alpha, \nu)x^3 + D(\alpha, \nu)x^{3+\alpha} + E(\alpha, \nu)]. \quad \text{(A5)}$$

Plugging these solutions into the governing equations, and matching terms of similar order in *x* for each equation, we obtain a linear system of algebraic equations for the unknown coefficients

$$A2,\ x^2 \to 4A - C = 1, \quad \text{(A6)}$$

$$A2,\ x^{2+\alpha} \to (4 + \alpha)B - D = -1, \quad \text{(A7)}$$

$$A2,\ x^{-1} \to E = -(A + B), \quad \text{(A8)}$$

$$A3,\quad x^2 \to 3A + 3C = (1 + \nu), \quad \text{(A9)}$$

$$A3,\quad x^{2+\alpha} \to (3 + \alpha)B + (3 + \alpha)D = -(1 + \nu). \quad \text{(A10)}$$

The solution of this system of equations is

$$A(\alpha, \nu) = F(0, \nu), \quad \text{(A11)}$$

$$B(\alpha, \nu) = -F(\alpha, \nu), \quad \text{(A12)}$$

$$C(\alpha, \nu) = G(0, \nu), \quad \text{(A13)}$$

$$D(\alpha, \nu) = -G(\alpha, \nu), \quad \text{(A14)}$$

$$E(\alpha, \nu) = -F(0, \nu) + F(\alpha, \nu), \quad \text{(A15)}$$

where

$$\boldsymbol{F}(\boldsymbol{\alpha},\boldsymbol{\nu}) = \frac{4+\alpha+\nu}{(3+\alpha)(5+\alpha)}, \qquad \text{(A16)}$$

$$\boldsymbol{G}(\boldsymbol{\alpha},\boldsymbol{\nu}) = (\mathbf{4}+\boldsymbol{\alpha})\boldsymbol{F}(\boldsymbol{\alpha},\boldsymbol{\nu}) - \mathbf{1}. \qquad \text{(A17)}$$

Bending Problem: We validated our procedure to recover monolayer tension caused by bending for a cell monolayer island under the normal traction stress:

$$\boldsymbol{T_z} = \boldsymbol{T_0}\boldsymbol{x}^2(\mathbf{1}-\boldsymbol{x}^{\alpha})\left(\boldsymbol{x}^{\alpha} - \frac{2}{2+\alpha}\right), \qquad \text{(A18)}$$

where *r* is the distance to the center of the island, *R* is the island radius and $x = r/R$. The parameter $\alpha \geq 1$ defines how concentrated the traction stresses are near the edge of the island, similar to the lateral validation problem, and $T_0$ has dimensions of stress. This synthetic traction stress distribution is in equilibrium of forces and moments, particularly bending moments. The governing equation for this axially symmetric problem is

$$\boldsymbol{\partial_{rr}M} + \frac{1}{r}\boldsymbol{\partial_r M} = \boldsymbol{T_z}. \qquad \text{(A19)}$$

Considering the forms of this equation and the forcing term, we try a solution of the form

$$\boldsymbol{M} = \boldsymbol{T_0}\boldsymbol{R}^2\left[\boldsymbol{A}(\boldsymbol{\alpha})\boldsymbol{x}^{\alpha+4} + \boldsymbol{B}(\boldsymbol{\alpha})\boldsymbol{x}^{2\alpha+4} + \boldsymbol{C}(\boldsymbol{\alpha})\boldsymbol{x}^{4}\right]. \qquad \text{(A20)}$$

Note that this solution automatically satisfies the boundary condition $\partial_r M|_{r=R} = 0$. Plugging this solution into the governing equation and matching terms of similar order in *x,* we arrive at:

$$\boldsymbol{x}^{2+\alpha} \to \boldsymbol{A} = \frac{1}{(2+\alpha)(4+\alpha)}, \qquad \text{(A21)}$$

$$\boldsymbol{x}^{2+2\alpha} \to \boldsymbol{B} = -\frac{1}{4(2+\alpha)^2}, \qquad \text{(A22)}$$

$$\boldsymbol{x}^{2} \to \boldsymbol{C} = -\frac{1}{8(2+\alpha)}. \qquad \text{(A23)}$$

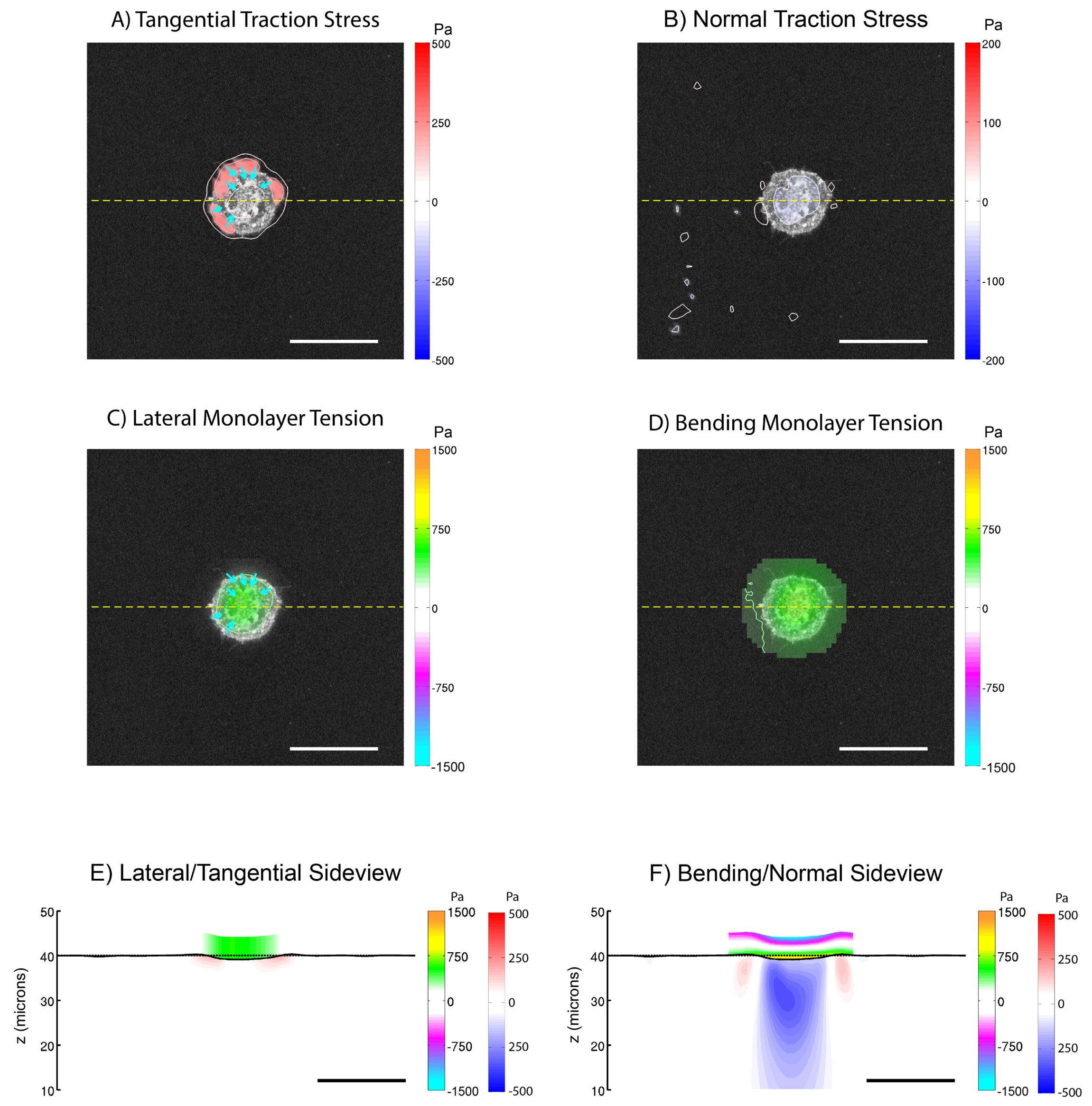


**Figure SM 1**

(A) Tangential traction stress $(\tau_{xz}, \tau_{yz})$ field on a 25-micron-radius cell island. The colormap represents magnitude of $\sqrt{\tau_{xz}^2 + \tau_{yz}^2}$ . (B) Normal traction stress field, with positive values (red) pointing upwards and away from the substrate. (C) Intracellular tension due to lateral deformation (color), with tangential traction stress (arrows) superimposed for reference. (D) Intracellular tension due to bending stress. (E) Side view representation of the intracellular and substrate stresses caused by lateral deformation along a representative line (yellow-dashed lines in A, B, C, and D). The black solid line represents the free surface of the deformed substrate. The red-blue colormap represents the magnitude of $\sqrt{\tau_{xz}^2 + \tau_{yz}^2}$ inside the gel. The orange-cyan colormap represents the intracellular tension. (F) Side view representing the intracellular tension due to bending stresses. Red-blue colormap represents the magnitude of $\tau_{zz}$ inside the gel. The orange-cyan colormap represents the intracellular tension due to bending. Scalebar = 50 μm.

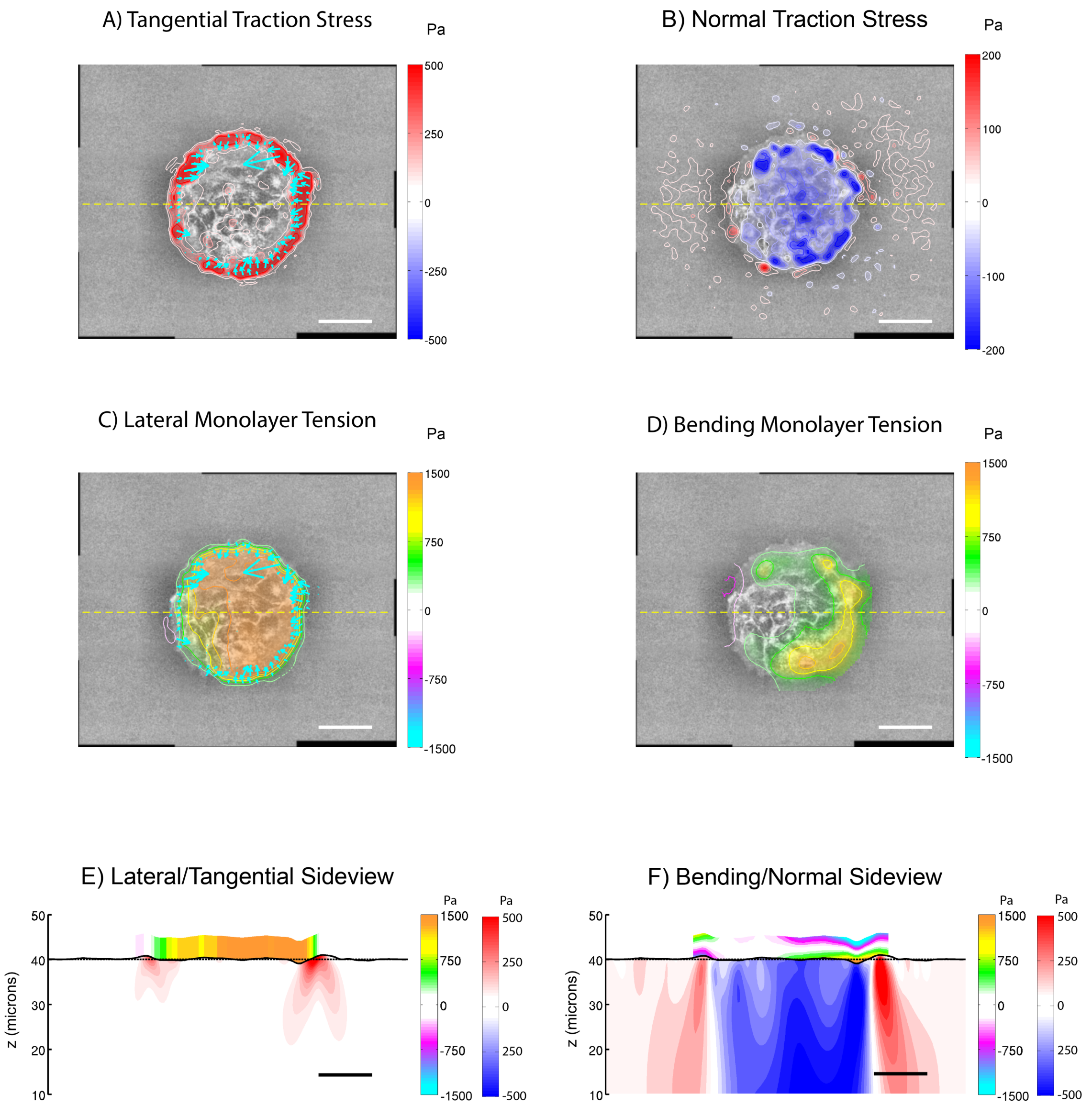


**Figure SM 2**

(A) Tangential traction stress $(\tau_{xz}, \tau_{yz})$ field on a 65-micron-radius cell island. The colormap represents magnitude of $\sqrt{\tau_{xz}^2 + \tau_{yz}^2}$ . (B) Normal traction stress field, with positive values (red) pointing upwards and away from the substrate. (C) Intracellular tension due to lateral deformation (color), with tangential traction stress (arrows) superimposed for reference. (D) Intracellular tension due to bending stress. (E) Side view representation of the intracellular and substrate stresses caused by lateral deformation along a representative line (yellow-dashed lines in A, B, C, and D). The black solid line represents the free surface of the deformed substrate. The red-blue colormap represents the magnitude of $\sqrt{\tau_{xz}^2 + \tau_{yz}^2}$ inside the gel. The orange-cyan colormap represents the intracellular tension. (F) Side view representing the intracellular tension due to bending stresses. Red-blue colormap represents the magnitude of $\tau_{zz}$ inside the gel. The orange-cyan colormap represents the intracellular tension due to bending. Scalebar = 50 μm.

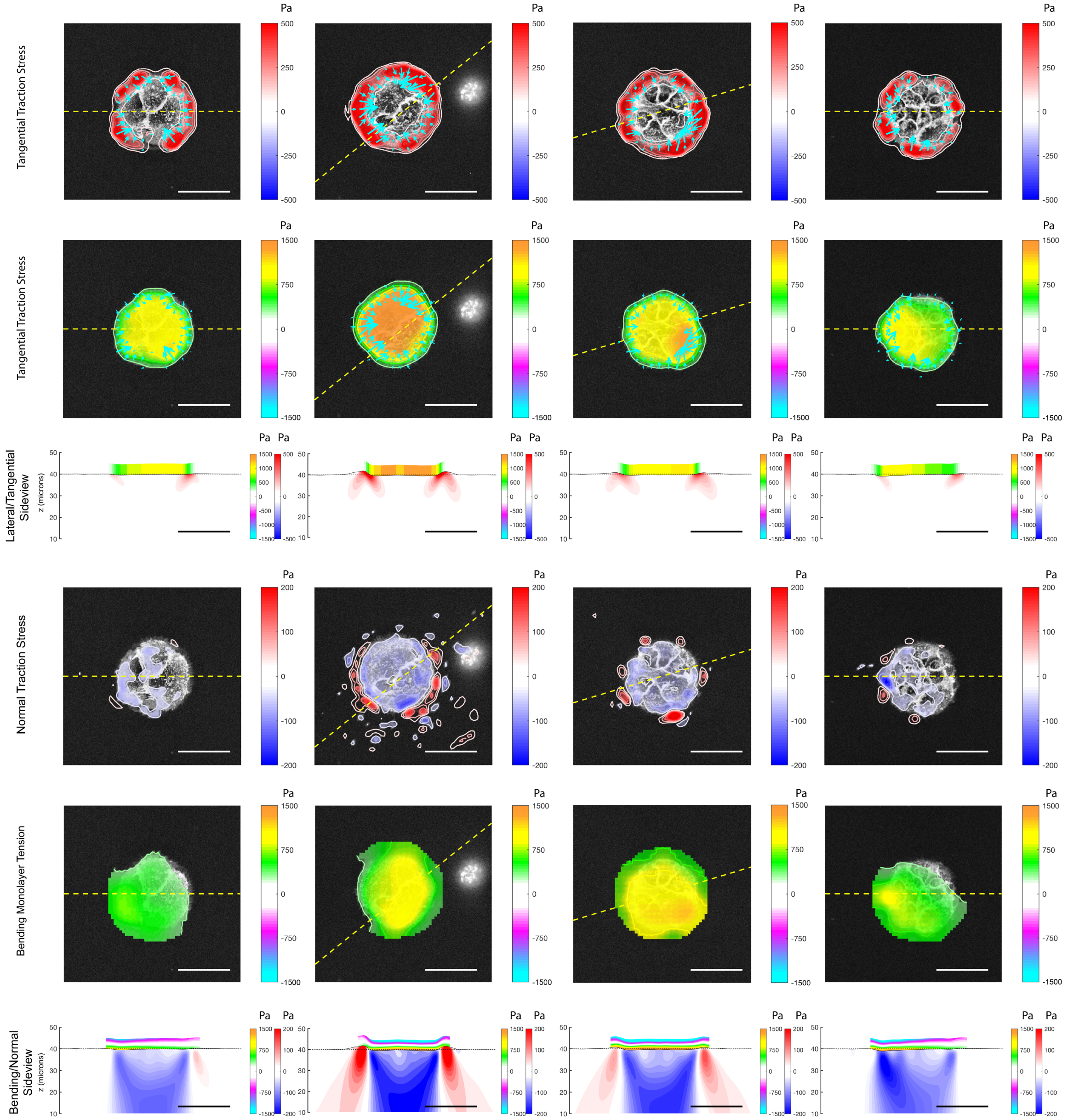


### Figure SM 3

Montage of traction stress and intracellular tension for N=4 different circular islands of 45 μm diameter. For each island, the data are represented as in Figure 5 of the main manuscript.

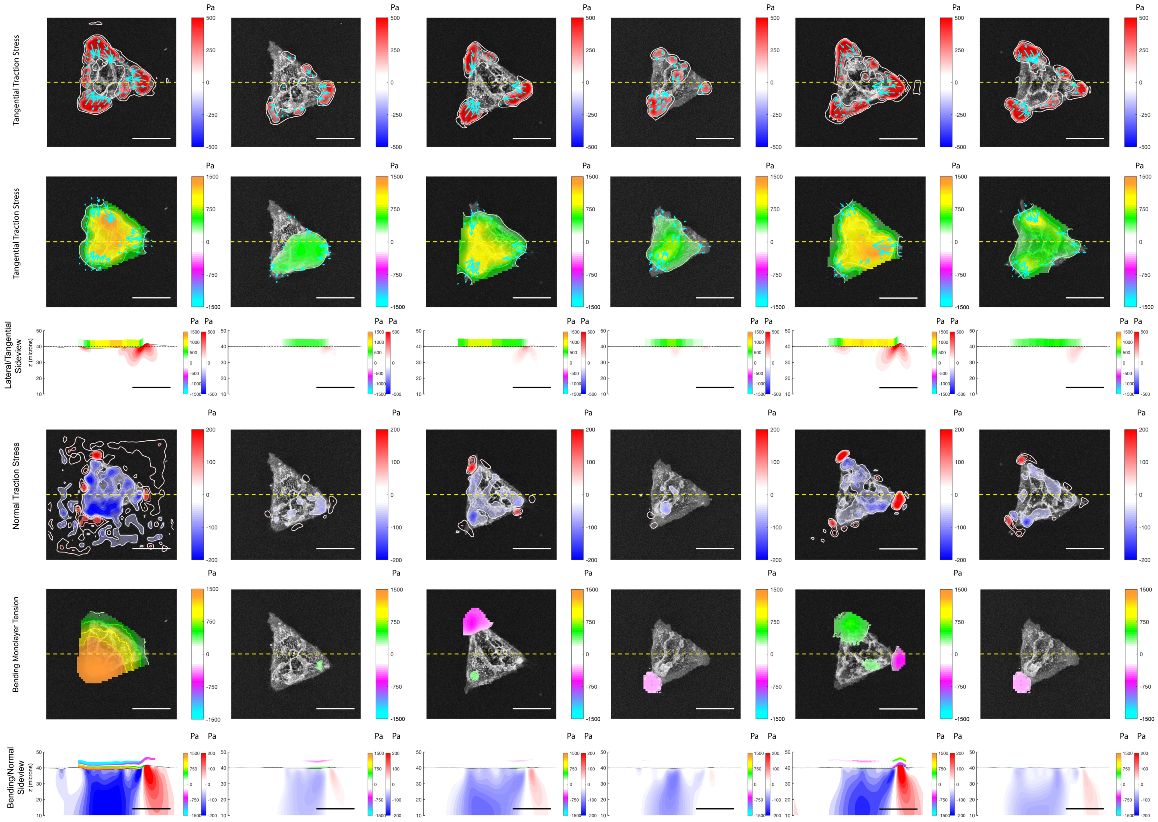


**Figure SM 4**

Montage of traction stress and intracellular tension for N=6 different triangular islands of 100 μm side. For each island, the data are represented as in Figure 6 of the main manuscript.

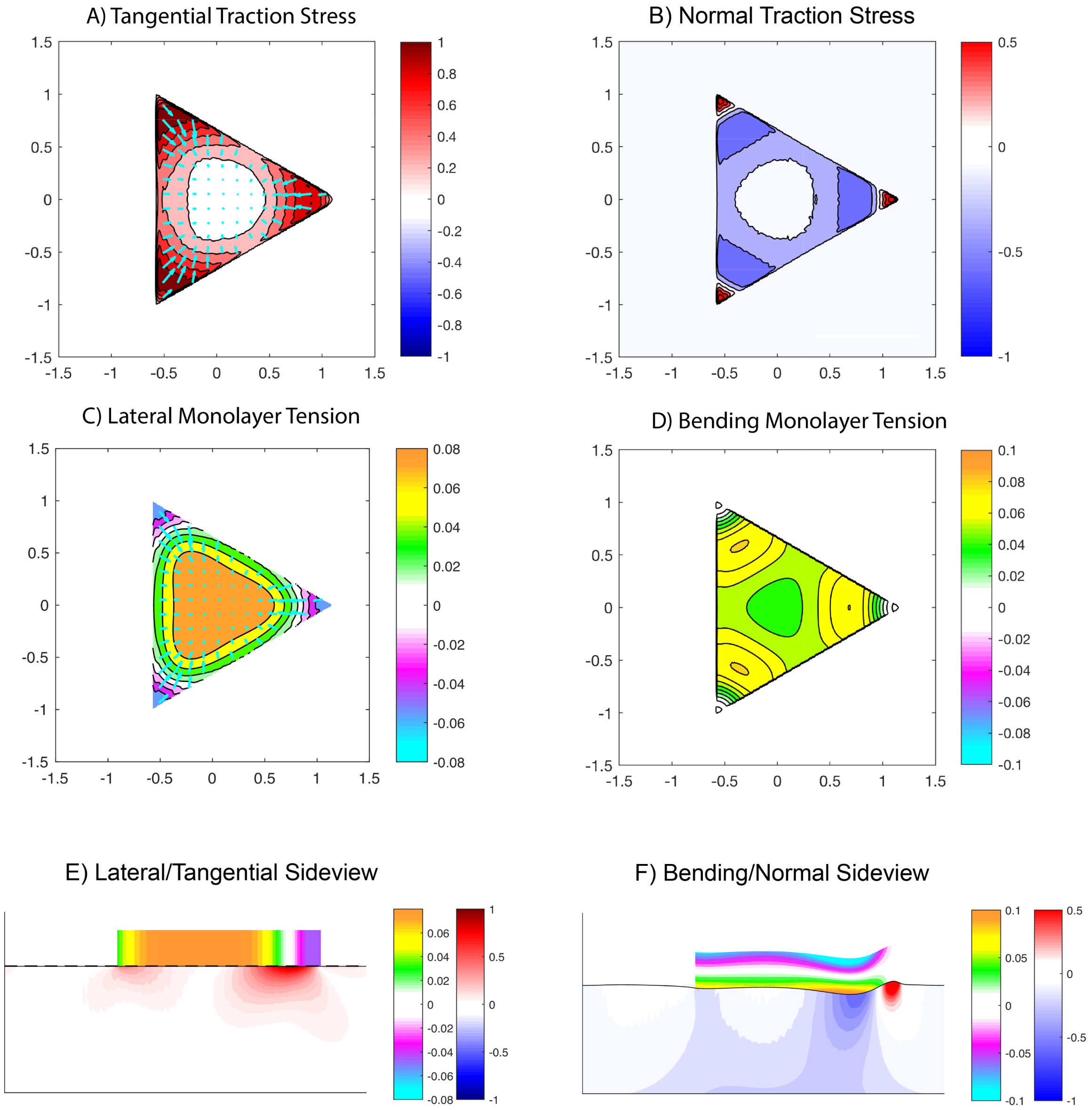


**Figure SM 5**

(A) Synthetic tangential traction stress $(\tau_{xz}, \tau_{yz})$ field on a simulated triangular geometry. The colormap represents magnitude of $\sqrt{\tau_{xz}^2 + \tau_{yz}^2}$. (B) Normal traction stress field, with positive values (red) pointing upwards and away from the substrate. (C) Intracellular tension due to lateral deformation (color), with tangential traction stress (arrows) superimposed for reference. (D) Intracellular tension due to bending stress. (E) Side view representation of the intracellular and substrate stresses caused by lateral deformation along a representative line (yellow-dashed lines in A, B, C, and D). The black solid line represents the free surface of the deformed substrate. The red-blue colormap represents the magnitude of $\sqrt{\tau_{xz}^2 + \tau_{yz}^2}$ inside the gel. The orange-cyan colormap represents the intracellular tension. (F) Side view representing the intracellular tension due to bending stresses. Red-blue colormap represents the magnitude of $\tau_{zz}$ inside the gel. The orange-cyan colormap represents the intracellular tension due to bending.

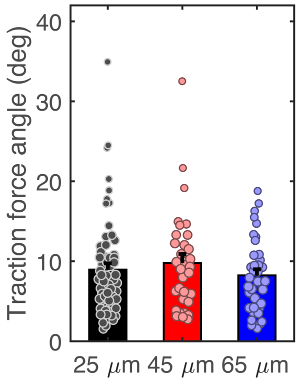


**Figure SM 6**

(A) Scatter plot of the peak tangential and normal traction stresses along radial lines of each circular island equally-spaced by a 45-degree angle. The color indicates the radius of island from which each point originates: black 25μm, red 45μm, and blue 65μm. (B) Angle at which cellular traction stresses pull upward and away from the substrate, estimated as the arctangent of the ratio between the peak normal and tangential traction stress for each radial line sampled. N = 64, 32, and 40 for the 25-, 45- and 65-μm-radius islands (8 data-points per cell island).

| $d_{edge}$(μm) | 0 | 10 | 20 | 40 |
|---|---|---|---|---|
| $\|\tau_{tan}\|$ | 0.16 | 0.24 | 0.03 | 0.02 |
| $\tau_{zz}$ | 0.15 | 0.24 | 0.66 | 0.06 |
| $s_{lat}$ | 0.20 | 0.18 | 0.22 | 0.29 |
| $s_{ben}$ | 0.14 | 0.28 | 0.43 | 0.06 |
| $\rho_{ben\text{-}lat}$ | 0.04 | 0.20 | 0.16 | 0.02 |

**Table SM 1**
P-values result of a Kruskal-Wallis test comparing the different average profiles shown in Figure 7 at 0, 10, 20, and 40 μm away from the edge of the island. $|\tau_{tan}|$ tangential traction stresses. $\tau_{zz}$ normal traction stresses. $s_{lat}$ lateral tension. $s_{ben}$ bending tension. $\rho_{ben\text{-}lat}$ relative contribution of lateral versus bending tension (see eq. 40 in the main text).

| $d_{edge}$(μm) | 0 | 10 | 20 | 40 |
|---|---|---|---|---|
| $\|\tau_{tan}\|$ | 0.12 | 0.04 | 0.01 | 0.02 |
| $\tau_{zz}$ | 0.03 | 0.006 | 0.006 | 0.02 |
| $s_{lat}$ | 0.01 | 0.005 | 0.006 | 0.02 |
| $s_{ben}$ | 0.002 | 0.002 | 0.002 | 0.02 |

**Table SM 2**
P-values result of a Kruskal-Wallis test comparing the different standard deviation profiles shown in Figure 7 at 0, 10, 20, and 40 μm away from the edge of the island.
